# Highly plastic macrophage niches orchestrate acquired quiescence and reactivation in breast-cancer bone metastasis

**DOI:** 10.64898/2026.08.31.748298

**Authors:** Yasuto Takeuchi, Huazi Zhang, Takahiko Murayama, Tsunaki Hongu, Kazuhiro Ikeda, Kuniko Horie, Satoshi Inoue, Masao Yano, Masahiko Tanabe, Yuki Kurokawa, Hirofumi Terakawa, Noriyuki Inaki, Kei-ichiro Tada, Etsuo A. Susaki, Eishu Hirata, Shu-ichi Tsukamoto, Masafumi Horie, Hiroshi Watarai, Kazuo Okamoto, Shinya Sato, Yohei Miyagi, Fumio Arai, Toshio Suda, Arinobu Tojo, Koji Okamoto, Hans Clevers, Tuomas Tammela, Noriko Gotoh

## Abstract

Recurrence and metastasis remain major causes of cancer mortality, sustained by therapy-resistant micrometastatic cells. Bone is a frequent site of breast-cancer relapse, yet the cues that reawaken disseminated cells remain poorly defined. We identify a previously unrecognized, highly plastic CXCL16⁺ macrophage population that integrates tumor-associated macrophage programs found in distant metastatic sites such as lung and brain with non-tumor disease–associated traits in bone marrow. These CXCL16⁺ macrophages establish a transient niche that restrains disseminated cancer-cell proliferation. Single-cell transcriptomics delineate functional remodeling of myeloid niches within the bone metastatic microenvironment: a CXCL16⁺ macrophage niche that transiently constrains metastatic growth, and G-CSF⁺ macrophage and neutrophil niches that reignite tumor outgrowth. In primary tumors, cancer-associated fibroblasts (CAFs) aberrantly secrete G-CSF in response to cancer-cell signals, expanding G-CSF-receptor–positive subset of cancer cells with high metastatic potential. In advanced human bone metastases, CXCL16⁺ macrophages localize to CAF-rich stroma but are excluded from cancer-cell clusters, indicating immune evasion. Together, these findings uncover CAF–bone-marrow cross-talk as a therapeutic target linking stromal inflammation, immune remodeling, and metastatic progression.

## Introduction

Despite advances in targeted and immune therapies, metastasis and recurrence remain the leading causes of cancer mortality^1^. Bone is one of the most frequent sites of relapse in breast cancer^2–4^, yet the mechanisms governing metastatic initiation in the bone marrow remain poorly defined. No effective targeted therapy exists to prevent or eradicate skeletal metastasis.

Breast cancer cells can disseminate early during tumor progression^5^. Disseminated tumor cells (DTCs) persist in a dormant state within specialized microenvironments of the bone marrow, including the perivascular and periosteal/osteogenic niches^6^. Under physiological conditions, these niches maintain hematopoietic stem cells (HSCs) in quiescence^7–9^. DTCs compete with HSCs for lodging within the perivascular niche by engaging endothelial cells and extracellular matrix components^10–13^, while interactions with osteogenic lineage cells, such as osteoblasts and bone-lining cells, anchor them to the periosteal surface^13, 14^. Perturbations of this equilibrium—through osteoblast-osteoclast dynamics or matrix remodeling—can trigger metastatic outgrowth, a “vicious cycle” of osteolysis and tumor proliferation. Clinically, anti-resorptive agents such as bisphosphonates slow skeletal complications but rarely eradicate metastatic disease, underscoring the need for mechanistic insight into the pathophysiology of bone metastasis^1, 6^.

The bone marrow is densely populated with hematopoietic cells spanning multiple lineages, yet how DTCs remodel or exploit this complex ecosystem to initiate metastasis remains largely unknown. Malignant cells in tumors are highly heterogeneous, comprising subpopulations with stem-like properties^15, 16^. These cells survive under non-adherent conditions, forming spheroids in vitro and resist anoikis—behaviors reminiscent of circulating DTCs. However, the molecular determinants linking stemness of cancer cells to bone-specific metastatic competence have remained elusive.

Recent single-cell omics technologies have illuminated the extraordinary heterogeneity of cancer cells, revealing trajectories that resemble developmental or regenerative processes^17–19^. Within this continuum, cancer stem-like cells can give rise to a drug-tolerant persister subpopulation^20^. Yet how the heterogeneity of bone-marrow– resident cells is remodeled upon invasion by DTCs remains largely unknown.

Under pathological conditions, bone-marrow–derived monocytes are recruited to inflamed tissues, where they differentiate into macrophages which play critical roles in cancer progression and in a multitude of other diseases besides cancer^21^. Historically, tumor-associated macrophages (TAMs)—the most abundant innate-immune population in tumors—have been divided into two polarization states: pro-inflammatory, anti-tumorigenic M1 macrophages that emerge early in tumorigenesis, and anti-inflammatory, pro-tumorigenic M2 macrophages that dominate later stages. However, recent single-cell transcriptomic analyses have revealed far greater complexity, with macrophage phenotypes varying profoundly across cancer types, organs, and even sub-anatomical microenvironments.

Triple-negative breast cancers (TNBCs), defined by the absence of estrogen receptor, progesterone receptor, and HER2 expression, exhibit an aggressive clinical course characterized by high rates of recurrence and metastasis^22, 23^. Among TNBCs, tumors with abundant desmoplastic stroma dominated by CAFs show particularly poor prognosis, underscoring the clinical relevance of stromal remodeling in this subtype^24, 25^.

Profiling of patient-derived CAF revealed aberrant G-CSF secretion, which induced G-CSF receptor (G-CSFR) expression and metastatic stem-like traits in cancer cells. Prophylactic G-CSF in vivo further amplified post-chemotherapy recurrence and bone metastasis, raising important clinical caution given its routine use to prevent chemotherapy-induced neutropenia^26^. Single-cell RNA sequencing (scRNA-seq) of bone marrow during early metastatic colonization uncovered a highly plastic CXC-chemokine-ligand-16– positive (CXCL16⁺) macrophage niche that restrains DTC proliferation, while coexisting G-CSF–responsive myeloid states open the gate for metastatic expansion. Human bone metastases show CAF-rich regions consistent with both microenvironments. Together, these findings unveil a CAF-driven G-CSF–G-CSFR circuit that rewires myeloid programs and forges a stromal–hematopoietic axis powering recurrence, bone metastasis, and therapy resistance.

## Results

### Patient-derived CAFs secrete G-CSF that induces stemness in breast cancer cells

To dissect the stromal factors that sustain stem-like properties, we established matched cultures of patient-derived breast cancer cells (BCCs) (BCC-P1–P4) and CAFs (CAF-P5–P9) from primary breast tumors (Fig. 1a; Extended Data Fig. 1a; Supplementary Table 1). Immunoblotting confirmed that BCCs retained the epithelial marker E-cadherin, whereas CAFs expressed fibroblast markers N-cadherin, vimentin, and α-SMA (Extended Data Fig. 1b).

**Fig. 1.**
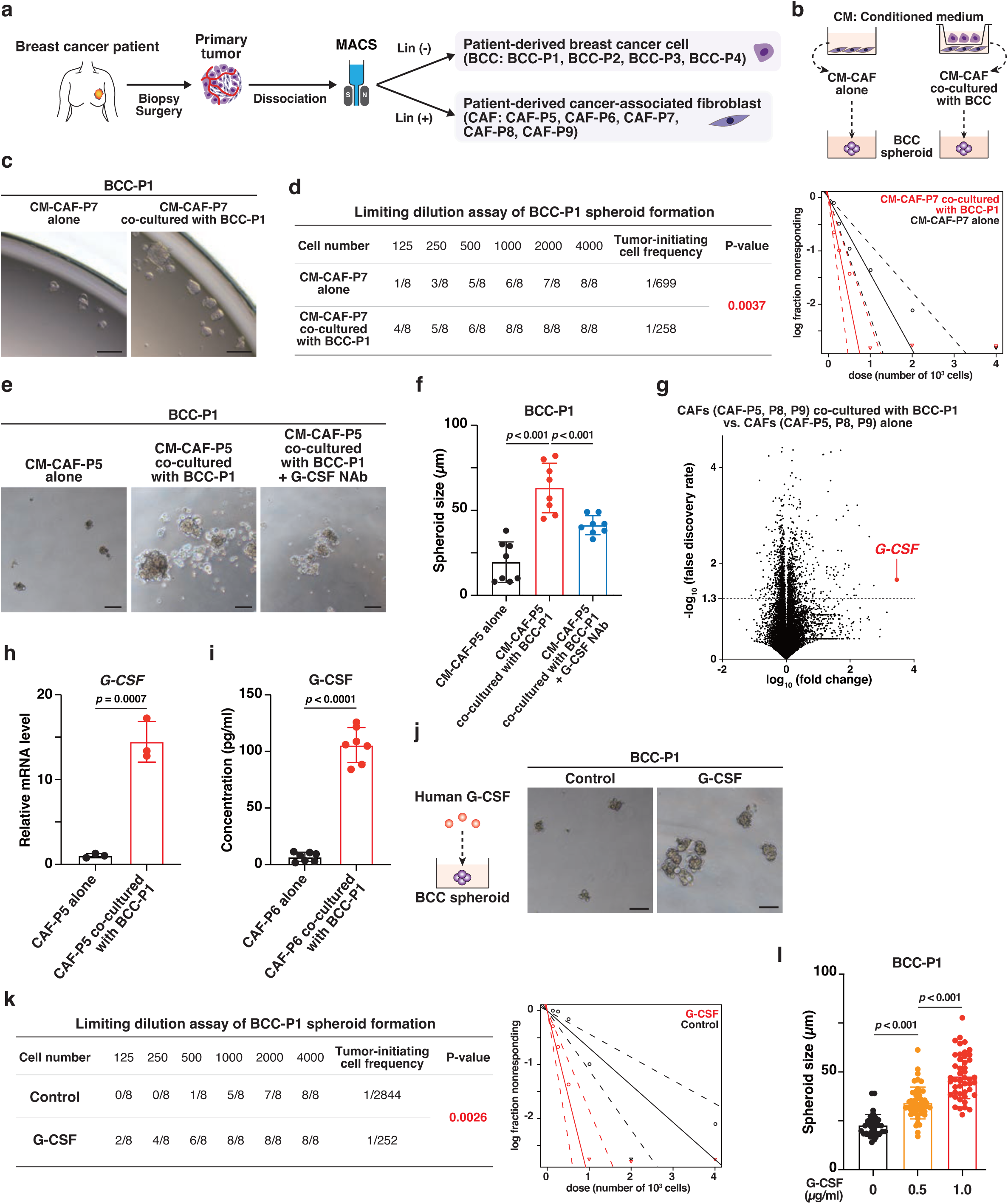
Patient-derived CAFs produce G-CSF to induce spheroid forming ability in patient-derived cancer cells. **a,** Schematic illustration of the workflow for establishing patient-derived breast cancer cells (BCCs) and cancer-associated fibroblasts (CAFs). **b,** Conditioned medium (CM) collected from CAFs cultured alone or from CAFs co-cultured with BCCs were used to treat BCC spheroids. **c, d,** Representative bright-field images (**c**) and extreme limiting dilution assay (ELDA) results (**d**) of BCC-P1 spheroids incubated with CM derived from CAF-P7 cultured alone or co-cultured with BCC-P1. Scale bars, 100 µm. **e, f,** Representative bright-field images (**e**) and quantification of spheroid size (**f**) of BCC-P1 spheroids incubated with the indicated CM. Scale bars, 100 µm. Statistical significance was determined by one-way ANOVA followed by post hoc multiple-comparisons test. Data represent mean ± SD (n = 8 spheroid cultures per group). Induction of tumor spheroid formation by CM derived from CAF-P5 co-cultured with BCC-P1 was attenuated by treatment with anti–G-CSF neutralizing antibodies (G-CSF NAb; 0.5 mg/ml). **g,** Volcano plot showing log₁₀ fold change in gene expression versus corresponding false discovery rate (FDR), comparing CAF-P5, CAF-P8, and CAF-P9 co-cultured with BCC-P1 versus those cultured alone (n = 3 biologically independent groups). **h, i,** Relative *G-CSF* mRNA levels (**h**) and secreted G-CSF protein levels (**i**) in CAFs cultured alone or co-cultured with BCC-P1, measured by qPCR and ELISA, respectively. Values were normalized to *GAPDH* or control CAFs. Statistical significance was determined by unpaired two-tailed Student’s *t*-test. Data are mean ± SD. **j–l,** Representative bright-field images (**j**), ELDA results (**k**), and quantification of spheroid size (**l**) of BCC-P1 spheroids cultured with or without recombinant G-CSF (0.5 or 1.0 µg/ml for 7 days). Scale bars, 100 µm. G-CSF induced tumor spheroid formation in a dose-dependent manner. Statistical significance was determined by one-way ANOVA with Dunnett’s post hoc test.

Conditioned medium (CM) from CAF–BCC co-cultures markedly increased tumor-spheroid number and size in both patient-derived BCCs and established cell lines BT-20 and BT-474 compared with CM from CAF monocultures (Fig. 1b–f; Extended Data Fig. 1c–f). Because spheroid formation reflects stemness activity, these data suggested that CAFs secrete paracrine factors enhancing stemness.

Bulk RNA-sequencing of CAFs cultured alone or co-cultured with BCCs revealed strong upregulation of *CSF3* (encoding G-CSF), ranking among the top induced transcripts (Fig. 1g; Supplementary Table 2). Quantitative PCR (qPCR) and enzyme-linked immunosorbent assay (ELISA) confirmed robust increases in *CSF3* mRNA and secreted G-CSF protein in co-cultured CAFs (Fig. 1h,i). Deconvolution of CAF transcriptomes using reference single-cell datasets^27^ showed that all three patient-derived CAF lines transcriptionally resembled tumor-like CAFs (tCAFs), a subtype localized near tumor–stroma borders (Extended Data Fig. 1g). tCAFs displayed the highest *G-CSF* expression among known CAF subsets, indicating that tumor-proximal fibroblasts are a major G-CSF source in breast cancer (Extended Data Fig. 1h).

Exogenous G-CSF treatment significantly increased spheroid-forming capacity in patient-derived BCCs and cell lines (Fig. 1j-l; Extended Data Fig. 1i), whereas a G-CSF-neutralizing antibody (G-CSF NAb) suppressed the CAF-induced enhancement (Fig. 1e,f). Gene-set enrichment analysis (GSEA) of RNA-seq data from co-cultured CAFs demonstrated activation of inflammatory transcriptional programs, including NF-κB signaling (Extended Data Fig. 1j). Immunofluorescence further revealed nuclear localization of NF-κB p65 in co-cultured CAFs but not in monoculture (Extended Data Fig. 1k,l), implicating NF-κB activation in G-CSF induction.

We next examined how BCCs stimulate NF-κB signaling in CAFs. RNA-seq of BCCs cultured with CAFs identified *CXCL12* as one of the most highly upregulated secreted factors (Extended Data Fig. 1m; Supplementary Table 3). Consistently, CAF co-culture increased *CXCL12* mRNA levels in BCCs, and recombinant CXCL12 induced *G-CSF* expression in CAFs (Extended Data Fig. 1n,o). These results define a paracrine loop in which CXCL12 secreted by BCCs activates NF-κB in CAFs, leading to G-CSF production that reinforces stemness properties in cancer cells.

### G-CSF–producing CAFs drive G-CSFR induction, tumor growth, and chemoresistant recurrence via the G-CSF–G-CSFR axis

G-CSF is known to induce its cognate receptor, G-CSFR, in responsive cells^28^. Consistently, co-culture of BCCs with patient-derived CAFs markedly upregulated *CSF3R* (encoding G-CSFR) mRNA and increased G-CSFR protein localization on the cancer-cell surface (Fig. 2a–c).

**Fig. 2.**
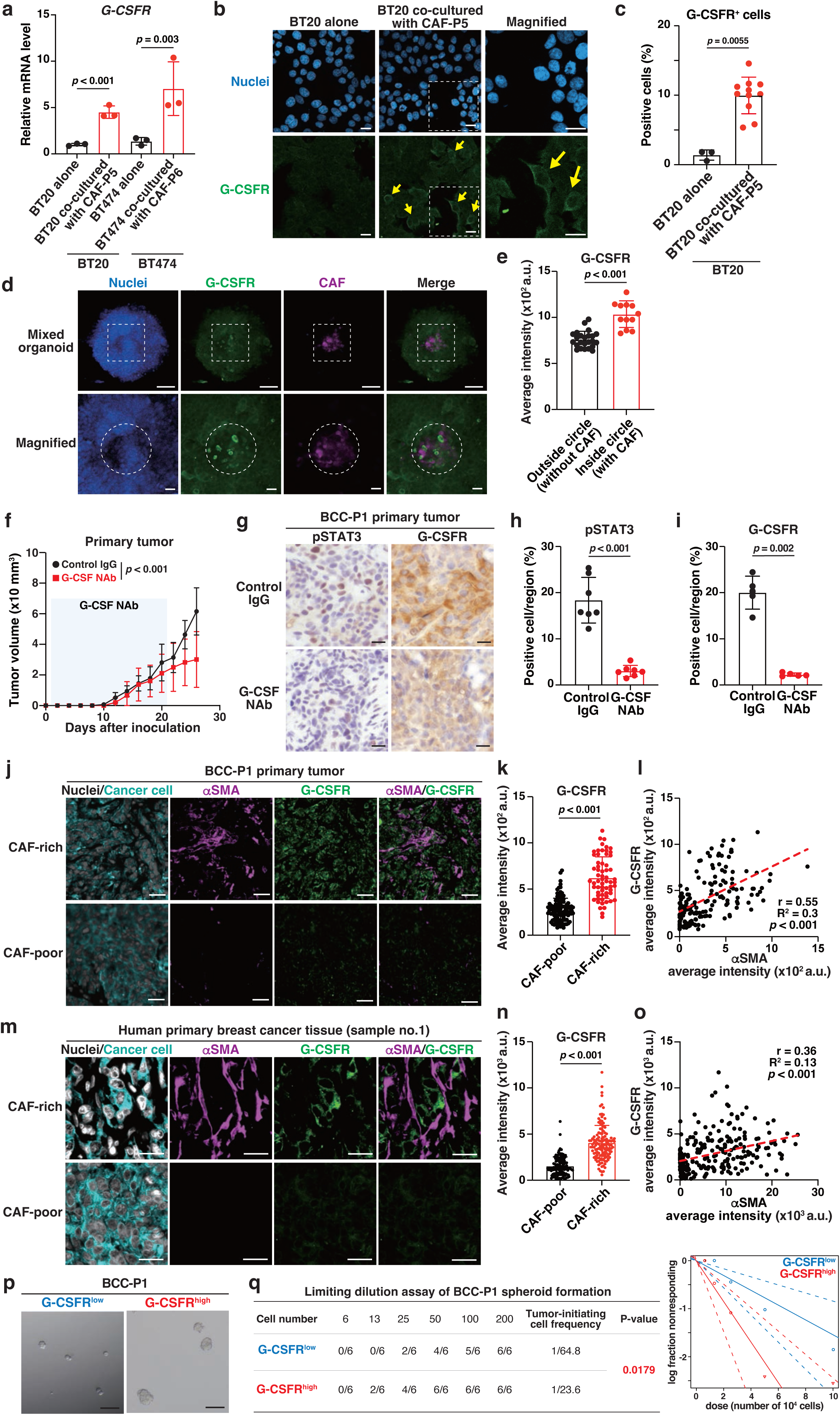
G-CSF-producing CAFs create a niche that induces G-CSFR expression and promotes tumor growth via the G-CSF–G-CSFR axis. **a,** Relative *G-CSFR* transcript levels in BT20 and BT474 BCCs cultured alone or co-cultured with CAF-P5 and CAF-P6, respectively (n = 3). **b,** Representative immunofluorescence images of G-CSFR in BT20 cells cultured alone or with CAF-P5. Arrows indicate plasma membrane–localized signals; magnified views of boxed areas are shown on the right. Scale bars, 100 µm. **c**, Quantification of G-CSFR^+^ BT20 cells cultured alone (n = 3 random fields) or with CAF-P5 (n = 11 random fields). **d,** Representative immunofluorescence images of G-CSFR in BCC-P1 organoids mixed with mCherry-labeled CAFs. Bottom panels show magnified boxed areas. Scale bars, 500 µm (top) and 100 µm (bottom). **e,** Quantification of G-CSFR signal intensity in CAF-containing (inside circle) and CAF-free (outside circle) regions in **d** (n = 12 and n = 24 random fields, respectively). **f,** Growth curves of PDX tumors treated with anti–G-CSF NAb (0.5 mg/kg, intraperitoneally once every two days). BCC-P1 cells were injected into the mammary fat pad. Data are mean ± SD (n = 3 mice per group). Statistical significance was determined by two-way ANOVA with post hoc multiple comparisons adjusted using the two-stage linear step-up procedure of Benjamini, Krieger and Yekutieli. **g,** Representative immunohistochemistry images of pSTAT3 and G-CSFR visualized by DAB (brown) with hematoxylin counterstain (purple). Scale bars, 100 µm. **h, i,** Quantification of signal intensities (n = 7 and 5 random fields, respectively). **j–o,** Representative immunofluorescence images of αSMA and G-CSFR in PDX tumors (**j**) and human breast cancer tissues (**m**) with quantifications (**k, n**) and Pearson correlation analyses (**l, o**). Scale bars, 100 µm. Data are mean ± SD. **p, q,** Representative bright-field images (**p**) and ELDA results (**q**) of BCC-P1 spheroids derived from G-CSFR^low^ and G-CSFR^high^ populations. Statistical significance in **a, c, e, h, i, k**, and **n** was determined by unpaired two-tailed Student’s *t*-test.

To mimic the tumor microenvironment in vitro, we established three-dimensional breast cancer organoids containing both CAFs and BCCs embedded in Matrigel^29^. Within these organoids, CAFs were positioned centrally and surrounded by cancer cells. Immunostaining revealed pronounced G-CSFR expression in cancer cells adjacent to CAF-rich regions (Fig. 2d,e), confirming that CAF-derived G-CSF induces G-CSFR expression in neighboring tumor cells.

To test the functional role of the G-CSF–G-CSFR axis in vivo, patient-derived xenograft (PDX) models were treated with G-CSF NAb. G-CSF blockade significantly suppressed primary tumor growth and reduced expression of phosphorylated STAT3 (pSTAT3), a downstream effector^30^, and G-CSFR in tumor tissues (Fig. 2f–i). Spatial analyses revealed strong correlations between CAF density, marked by α-SMA, and G-CSFR expression in both PDX tumors and human primary breast cancer samples (Fig. 2j–o; Extended Data Fig. 2a,b; Supplementary Table 4). These findings establish that G-CSF–producing tCAFs create a stromal niche that sustains G-CSFR⁺ cancer stem-like cells and promotes tumor expansion through the G-CSF–G-CSFR–STAT3 signaling axis.

We next assessed whether G-CSFR⁺ cells indeed possess stemness properties. Flow-cytometric sorting of BCC populations based on G-CSFR expression revealed that G-CSFR^high^ cells exhibited significantly greater spheroid-forming efficiency than G-CSFR^low^ cells in both patient-derived BCCs and BT-20 cell lines (Fig. 2p,q; Extended Data Fig. 2c-e), indicating enhanced stemness activity.

We treated patient-derived breast cancer organoids with clinically used-paclitaxel (PTX) alone or PTX in combination with exogenous G-CSF (Fig. 3a). Following PTX treatment, cell numbers were decreased, but the proportion of G-CSFR^high^ breast cancer stem-like cells increased and was further elevated by the addition of G-CSF (Fig. 3b,c; Extended Data Fig. 2f). The fraction of Ki67-positive cells within the G-CSFR^high^ population was also increased by G-CSF treatment (Fig. 3d,e). These findings posit that G-CSF may stimulate the proliferation of G-CSFR^high^ cancer stem-like cells that are resistant to chemotherapy in vitro.

**Fig. 3.**
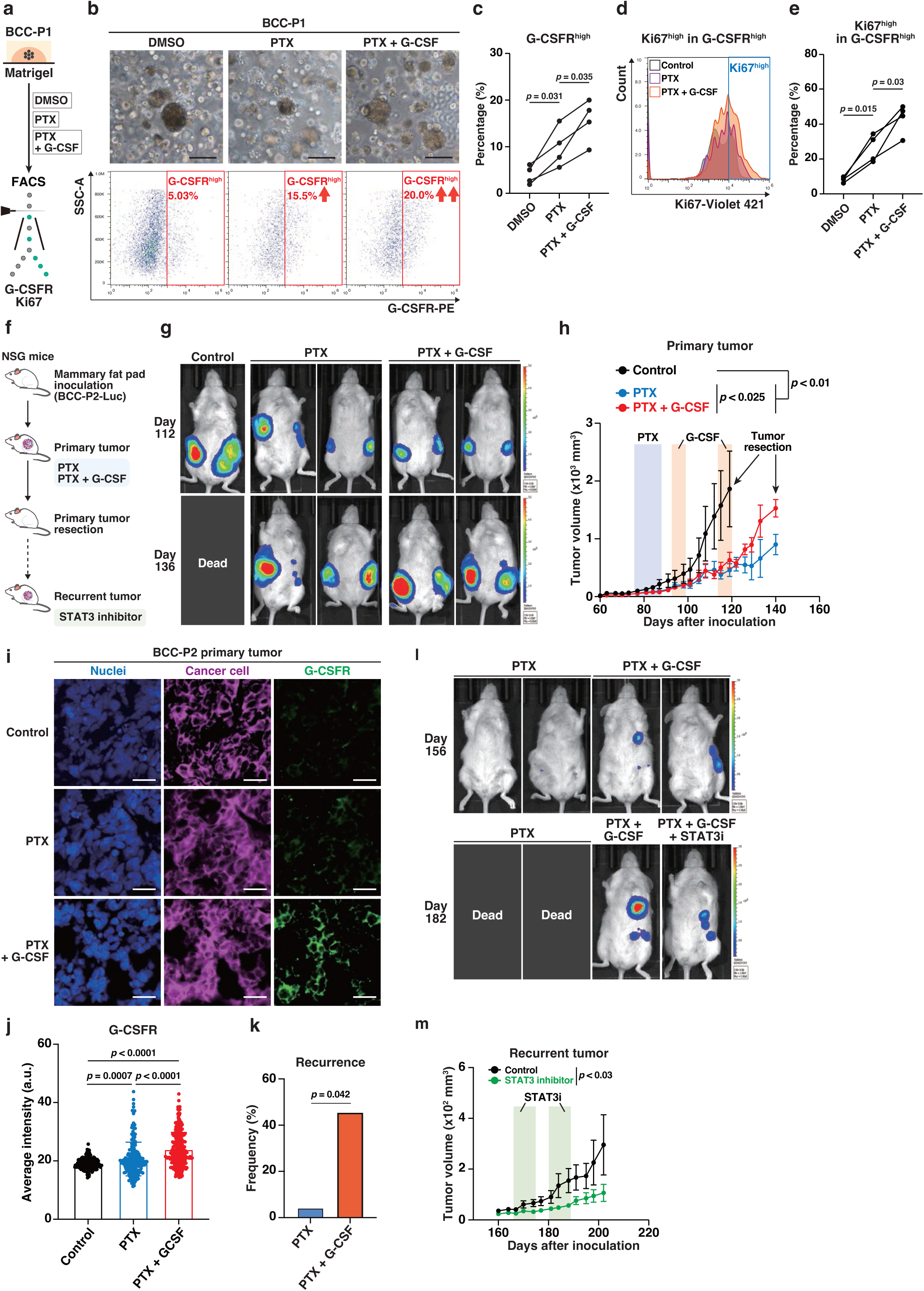
Exogenous G-CSF enhances chemoresistant G-CSFR^high^ cancer stem-like cells and promotes tumor regrowth and local recurrence. **a,** Schematic of the workflow for paclitaxel (PTX) (10 µM, 5 days) treatment with or without G-CSF (10 ng/ml for 3 days concurrently) in BCC-P1 organoids. **b,** Representative bright-field images of organoids (top) and flow cytometric analysis of G-CSFR^high^ cells (bottom). Scale bars, 100 µm. **c,** Quantification of G-CSFR^high^ cells (n = 4 per condition). **d, e,** Ki67^high^ cells within G-CSFR^high^ populations analyzed by flow cytometry (**d**) and quantification (**e**; n = 4 per condition). **f,** Schematic of in vivo treatment. BCC-P2-luciferase (BCC-P2–Luc) cells were implanted orthotopically and treated with PTX alone (10 mg/kg, intraperitoneally twice weekly for 2 weeks) or followed by G-CSF (125 µg/kg, intraperitoneally once daily for 5 days, for two consecutive cycles). **g,** Longitudinal IVIS bioluminescence imaging of tumorigenesis. **h,** Tumor growth curves of PDX models. Tumors were resected at ∼2,000 mm³ (control) or 140 days post-inoculation (treatment), after which local recurrence was monitored. Data represent mean ± SEM (n = 4 for control, 4 for PTX, and 8 for PTX + G-CSF). **i, j,** Representative immunofluorescence images of human cancer cell (STEM121; human-specific) and G-CSFR in BCC-P2 primary tumors (**i**) and quantification of G-CSFR fluorescence (**j;** n = 250, 207, and 269 fields for control, PTX, and PTX + G-CSF, respectively). Scale bars, 100 µm. Statistical significance was determined by one-way ANOVA with Dunnett’s post hoc test. **k,** Frequency of local recurrence analyzed by Fisher’s exact test. **l,** Longitudinal IVIS imaging of local recurrence. **m,** Recurrent tumor growth curves of PDX models treated with or without STAT3 inhibitor (100 mg/kg, intraperitoneally once daily for 1 week, for two consecutive cycles). Data represent mean ± SEM. **h, m**, Statistical significance was determined by two-way ANOVA with post hoc multiple comparisons adjusted using the two-stage linear step-up procedure of Benjamini, Krieger and Yekutieli.

To evaluate the clinical implication of these findings, we modeled neoadjuvant chemotherapy (NAC) in PDX-bearing mice using PTX with or without exogenous G-CSF administration. While PTX initially reduced tumor burden, combined treatment with G-CSF led to regrowth at markedly higher rates than PTX alone (Fig. 3f-h). Immunohistochemistry demonstrated increased fractions of G-CSFR^high^ tumor cells (Fig. 3i,j).

Longitudinal monitoring of tumor-bearing mice revealed that the addition of G-CSF accelerated recurrence after surgical resection of primary tumors (Fig. 3k,l). Importantly, treatment of recurrent tumors with a STAT3 inhibitor (STAT3i) significantly reduced recurrent growth, mirroring the effect of G-CSF neutralization (Fig. 3l,m).

Analysis of public breast cancer datasets^31^ showed that high *GCSF* expression correlates with an increased risk of relapse (Extended Data Fig. 2g). Together, these findings indicate that G-CSF given prophylactically or therapeutically during chemotherapy to mitigate infectious risk may unintentionally sustain and expand G-CSFR^high^ cancer stem-like cells, fostering tumor regrowth and local recurrence. Disrupting the G-CSF–G-CSFR axis thus represents a rational strategy to suppress tumor persistence and recurrence.

### G-CSFR⁺ cells act as bone-metastasis–initiating cells and remodel the bone-marrow microenvironment

Because G-CSF is physiologically abundant in bone marrow^30^, we hypothesized that G-CSFR^high^ cancer cells might preferentially colonize this niche. To test this, we injected sorted G-CSFR^high^ or G-CSFR^low^ populations into the caudal artery of immunodeficient mice to generate bone-metastatic PDX models (Extended Data Fig. 3a). G-CSFR^high^ cells produced significantly higher bone-metastatic burden than G-CSFR^low^ cells, confirming that the G-CSFR^high^ fraction contains metastasis-initiating cells (Fig. 4a–c).

**Fig. 4.**
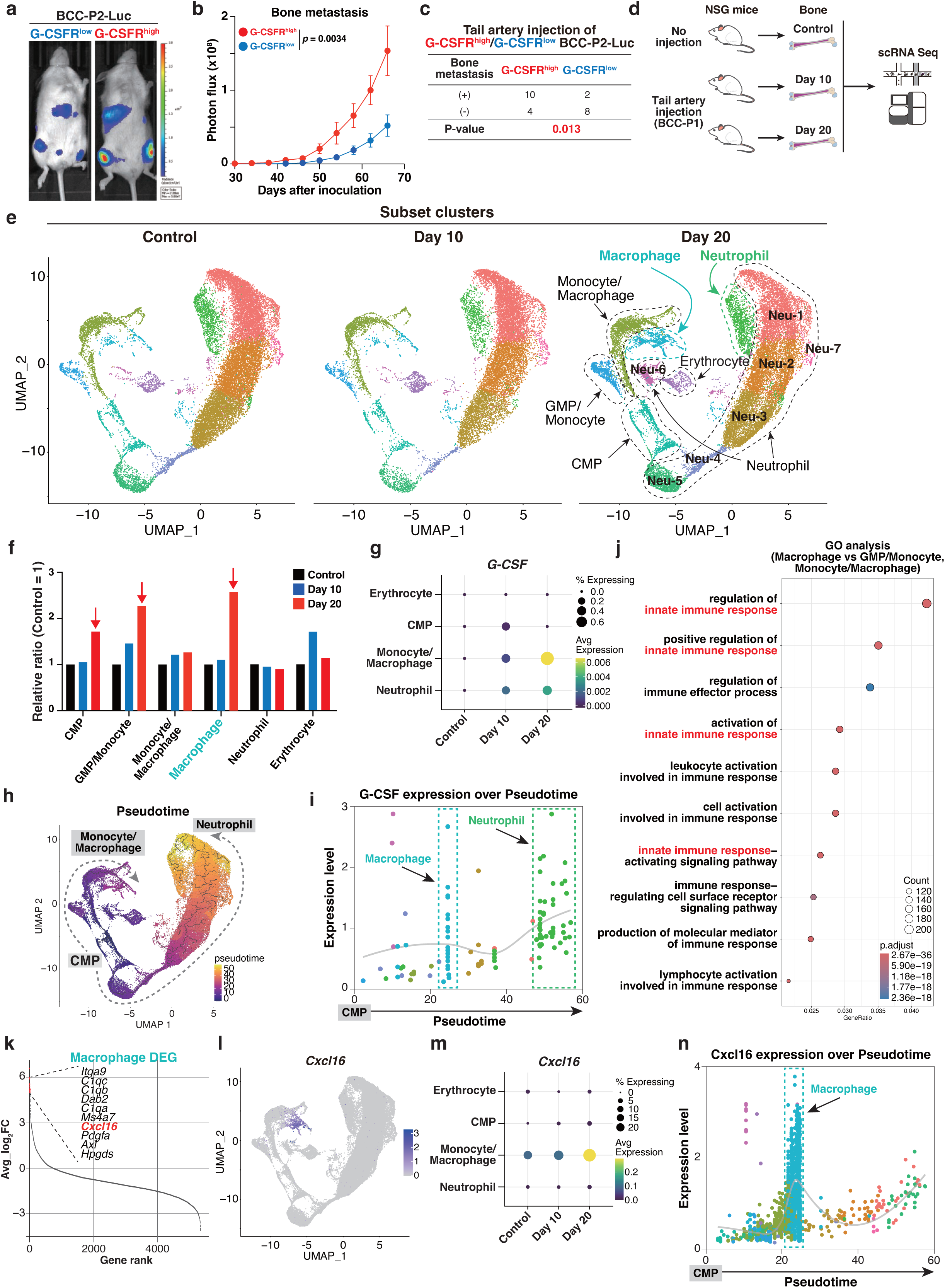
G-CSFR^high^ cells initiate bone metastasis, accompanied by expansion of CXCL16-expressing macrophages during metastatic onset. **a,** Representative IVIS bioluminescence images showing bone metastases at day 66. BCC-P2–Luc cells were injected into the caudal artery to generate bone metastasis. **b,** Quantification of bone metastasis by IVIS bioluminescence intensity. Statistical significance was determined by two-way ANOVA with post hoc multiple comparisons adjusted using the two-stage linear step-up procedure of Benjamini, Krieger and Yekutieli. Data represent mean ± SEM (n = 14 for G-CSFR^high^ and n = 10 for G-CSFR^low^). **c,** Incidence of bone metastasis. Statistical significance was determined by Fisher’s exact test. **d,** Schematic illustration of the workflow for single-cell RNA sequencing (scRNA-seq) during bone metastasis initiation. **e,** Uniform manifold approximation and projection (UMAP) visualization of re-clustered myeloid-lineage cells subset from bone metastasis–initiating lesions, colored by subclusters. **f,** Relative ratio of each myeloid subset normalized to corresponding control values. **g,** Dot plot showing *G-CSF* expression across cell populations over time; color indicates scaled mean expression and dot size indicates the fraction of expressing cells. **h,** Pseudotime trajectory of myeloid cells, with the common myeloid progenitor (CMP) cluster set as the root. Ridge plots show density estimates of each subcluster across pseudotime. Branching points denote inferred fate decisions, and color gradients indicate progression along the pseudotime axis. i, *G-CSF* expression dynamics along pseudotime. **j,** Representative Gene Ontology (GO) pathways enriched in the novel macrophage cluster compared with GMP/monocyte and monocyte/macrophage clusters. Color reflects adjusted *P* values; dot size indicates the number of genes per term. **k,** Differentially expressed genes (DEGs) in the novel macrophage cluster compared with all other clusters, ranked by average log₂ fold change. **l,** Feature plot of *Cxcl16* expression in UMAP visualization. **m,** Dot plot of *Cxcl16* expression across cell populations over time. n, *Cxcl16* expression dynamics along pseudotime.

At 20 days after injection, histological analyses revealed lesions ranging from micrometastases, composed of single DTCs, to early macrometastases forming compact clusters lacking CAF infiltration (Extended Data Fig. 3b). Approximately 10% of micrometastatic cancer cells were attached to CD31⁺ endothelial cells, whereas most were embedded among bone-marrow cells (Extended Data Fig. 3c,d), suggesting the existence of novel cellular niches beyond classical perivascular and periosteal compartments.

To capture the dynamic remodeling of the bone marrow microenvironment during metastatic progression, we isolated total bone marrow cells by conventional aspiration followed by mechanical fracture to encompass bone-lining populations at days 0, 10, and 20 after tail-artery injection and performed scRNA-seq (Fig. 4d). Unsupervised clustering spanned 12 clusters with distinct cell types and their proportions based on canonical marker gene expression (Extended Data Fig. 3e,f; Supplementary table 5). Majority were hematopoietic-lineage cells; neutrophil, monocyte/macrophage, erythrocytes, dendritic cells, basophil/eosinophils (Extended Data Fig. 3g). Although minor in abundance, chondrocyte/osteoblasts and endothelial cells underwent marked changes between day 10 and day 20 (Extended Data Fig. 3g,h). Thus interactions of metastatic BCCs with both periosteal and perivascular niches may take place, corroborating that our scRNA-seq data capture the initial phase of bone metastasis.

To improve resolution of the myeloid-lineage cell data, we next excluded relatively minor populations with defined cell lineages (dendritic cells, chondrocyte/osteoclasts, basophil/eosinophils and endothelial cells) and re-clustered the remaining populations. Unsupervised clustering identified 13 major cell populations (Fig. 4e; Extended Data Fig.4a; Supplementary Table 5). Although neutrophils were abundant (Extended Data Fig. 4b), the most pronounced temporal changes occurred in myeloid compartments, particularly common myeloid progenitors (CMPs), granulocyte–macrophage progenitors (GMPs), and macrophages (Fig. 4f; Extended Data Fig. 4b).

G-CSF expression (*Csf3*) was detected in CMP, monocyte/macrophage, and neutrophil populations and increased from day 10 to day 20 (Fig. 4g). Pseudotime trajectory analysis positioned CMP-like cells as the root of two major differentiation paths: a clockwise branch generating monocyte/macrophage lineages and a counterclockwise branch producing neutrophils, both associated with *G-CSF* upregulation (Fig. 4h,i; Extended Data Fig. 4c). These results suggest that macrophages and neutrophils are primary sources of G-CSF within the metastatic bone-marrow milieu, potentially supporting the survival and expansion of G-CSFR⁺ metastasis-initiating cells.

We next focused on macrophages positioned at the terminal end of the pseudotime trajectory, as their proportion expanded most markedly between day 10 and day 20 (Fig. 4e,f,h). Gene Ontology enrichment analysis revealed that these macrophages upregulated innate immune–response genes compared with other myeloid subsets (Fig. 4j). Among the top differentially expressed genes, *Cxcl16* ranked prominently (Fig. 4k; Supplementary table 6). Expression mapping confirmed that *Cxcl16* was specifically induced in macrophages and expanded over time (Fig. 4l-n), suggesting that *Cxcl16* marks a newly emergent macrophage population during early metastatic colonization.

### CXCL16^+^ macrophages orchestrate plasticity and heterogeneity during early bone metastasis

To characterize macrophage heterogeneity during the initial phase of bone metastasis, reclustering of the macrophage compartment identified eight transcriptionally distinct macrophage subclusters (Fig. 5a). Among these, cluster 1—distinguished by strong *Cxcl16* expression—expanded more than ten-fold between day 10 and day 20 (Fig. 5b,c; Extended Data Fig. 5a). Cluster 0 contained immature monocyte-derived macrophages expressing *Csf3* (G-CSF) (Extended Data Fig. 5b), and both clusters 0 and 1 showed high NF-κB-related gene-set enrichment, indicating inflammatory activation during early colonization (Extended Data Fig. 5c-e; Supplementary table 7). Pseudotime analysis delineated a transition from cluster 0 → 1, followed by bifurcation into clusters 2 and 3 (Fig. 5d,e). Overlay of *Cxcl16* expression revealed a sharp peak in cluster 1, moderate levels in clusters 2–4, and near absence in clusters 5–7 (Fig. 5c,f).

**Fig. 5.**
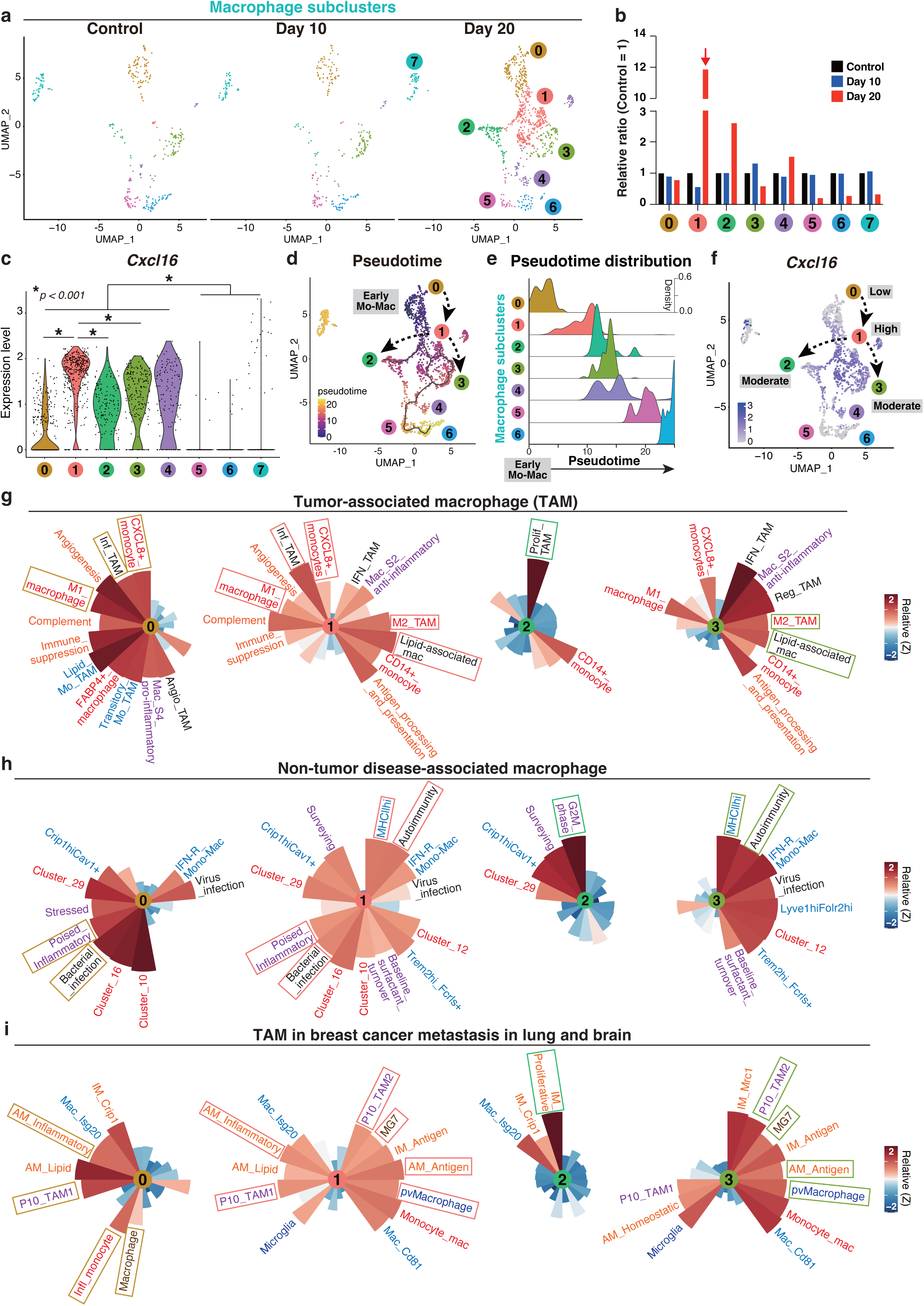
*Cxcl16* expression associates macrophage plasticity and contributes to their diversity. **a,** UMAP visualization of scRNA-seq data showing newly emerging macrophage clusters during the early phase of bone metastasis. **b,** Relative proportions of each macrophage subset compared with corresponding control samples. **c,** Violin plots showing *Cxcl16* expression across clusters in (**a**). Statistical significance was determined by one-way ANOVA with Bonferroni’s post hoc test. **d,** Pseudotime ordering of macrophage subclusters with the early monocyte-derived macrophage (Early Mo-Mac; cluster 0) set as the root. Cluster numbers indicate identities arranged along pseudotime progression. Ridge plots (thin lines) represents density estimates of cells from each subcluster along the pseudotime axis. **e,** Pseudotime distribution of macrophage subclusters. **f,** Feature plot showing *Cxcl16* expression mapped onto the UMAP. **g,** Rose plots showing relative AUCell-based enrichment of 19 tumor-associated macrophage (TAM) gene signatures across subclusters 0–3. Curated TAM gene sets are labeled in black, and publicly available signatures are shown in distinct colors corresponding to their respective references. Rose plots were generated from the cluster-averaged relative AUCell scores (heatmap of Extended Data Fig. 6), with the sector length scaled to Z-score-normalized values. **h,** Rose plots showing enrichment of 17 non-tumor disease-associated macrophage gene signatures across subclusters 0–3. Curated signatures are labeled in black, and publicly available ones are color-coded by source references. **i,** Rose plots showing enrichment of 20 macrophage gene signatures derived from TAMs in metastatic breast cancer (lung and brain) and from primary non-small-cell lung cancer (NSCLC) and glioma, across subclusters 0–3. Gene sets from PDX samples (P10-TAM1 and P10-TAM2) are labeled in purple, and other publicly available gene sets are shown in distinct colors corresponding to their respective references.

Because TAMs encompass diverse functional states, we compared each cluster with 19 curated and publicly available TAM signatures (Supplementary Tables 8)^21, 32–44^. Cluster 0 was enriched for inflammatory programs represented in the left side of the rose plots (CXCL8_monocyte, Inflammatory_TAM [Inf_TAM], M1_macrophage, and others) (left panel, Fig. 5g). In contrast, the CXCL16-high cluster 1 displayed a broader transcriptional diversity encompassing not only these inflammatory programs but also immune-regulatory signatures represented in the right side of the rose plots (M2_TAM, Lipid-associated_macrophage, and others) (second left panel, Fig. 5g). Cluster 2 adopted a more restricted, proliferative phenotype (Prolif_TAM), whereas cluster 3 exhibited dominant immune-regulatory programs (right panel, Fig. 5g).

The angiogenic TAM (Angio_TAM) signature was associated with cluster 0, reminiscent of perivascular niche formation (Extended Data Fig. 5g–i). In contrast, osteoclast-related genes were enriched in cluster 2, suggesting initiation of an osteoblast–osteoclast–tumor “vicious cycle” that sustains bone colonization (Extended Data Fig. 5j–l). These findings imply that CXCL16-high cluster 1 macrophages act as a bridge between perivascular (cluster 0) and osteogenic (cluster 2) niches.

We next examined whether CXCL16-expressing macrophages share transcriptional profiles with disease-associated macrophages identified in non-cancer contexts. Using 17 curated and publicly available gene sets derived from infection, bone fracture, osteoarthritis, autoimmunity, and senescence (Supplementary Table 9)^39, 45–64^, we found that acute inflammatory phenotypes were enriched in cluster 0 (Senescence-associated Poised_Inflammatory, Bacterial_infection, and others) (left panel, Fig. 5h), whereas chronic inflammatory phenotypes were enriched in cluster 3 (Osteoarthritis-associated [MHCIIhi], Autoimmunity, and others) (right panel, Fig. 5h). Both acute and chronic inflammatory phenotypes were broadly distributed across cluster 1, whereas proliferative signatures were confined to cluster 2 (G2M_phase) (center panels, Fig. 5h).

Furthermore, we interrogated whether CXCL16-expressing macrophages resemble TAMs found in breast cancer metastatic lesions of distant organs such as lung and brain. To this end, we used 20 publicly available gene sets together with those derived from our breast-cancer PDX model and its spontaneous lung metastases (Extended Data Fig. 8a,b)^65–67^, and included TAM signatures from primary tumors: non–small cell lung cancer (NSCLC) and glioma–for comparison^68, 69^ (Supplementary Table 10). We found that all the TAM profiles were represented in distinct clusters. Cluster 0 was enriched for inflammatory phenotypes, including lung metastasis–associated inflammatory alveolar macrophages (AM_Inflammatory), Patient 10 lung metastasis–associated TAM1 (P10_TAM1), NSCLC-associated inflammatory monocytes (Infl_monocyte), and glioma-associated macrophages (Macrophage) (left panel, Fig. 5i). In contrast, cluster 3 displayed immune-regulatory phenotypes, such as Patient 10 lung metastasis– associated TAM2 (P10_TAM2), glioma-associated MG7, lung metastasis–associated antigen-presenting alveolar macrophages (AM_Antigen), and brain metastasis– associated perivascular macrophages (pvMacrophage) (right panel, Fig. 5i). These distinct enrichment patterns indicate that cluster 0 macrophages adopt inflammatory and innate-immune activation programs, whereas cluster 3 macrophages acquire immune-regulatory and antigen-presentation traits typical of advanced metastatic niches. Notably, CXCL16-high cluster 1 macrophages encompassed signatures of both, underscoring their transitional and highly plastic nature bridging inflammatory and regulatory TAM states across metastatic sites (Fig. 5i).

Clustering analysis revealed consistent trends, with cluster 1 macrophages exhibiting broad transcriptional signatures (Extended Data Fig. 6a–c). Gene sets visualized on the left and right sides of the rose plots highlight similarities between tumor-associated macrophages (TAMs), non-tumor disease–associated macrophages, and TAMs identified in breast-cancer metastases to the lung and brain (Fig. 5g-i; Extended Data Fig. 7a-c).

Together, these analyses delineate CXCL16-expressing macrophages as highly plastic and heterogeneous populations that undergo dynamic differentiation along the monocyte–macrophage continuum during metastatic progression. Remarkably, CXCL16-expressing macrophages emerging during early bone metastasis already exhibit transcriptional features characteristic of macrophages found in advanced metastatic niches, including those in lung and brain.

### CXCL16⁺ macrophages transiently restrain, whereas G-CSF⁺ macrophages promote, metastatic outgrowth

Immunofluorescence of bone-metastatic PDX lesions revealed CXCL16⁺ F4/80⁺ macrophages encircling and occasionally contacting human cancer cells (Fig. 6a). Intriguingly, neutralization of CXCL16 accelerated bone-metastatic progression, indicating that these macrophages impose a transient proliferative restraint during early colonization (Fig. 6b,c). Consistently, Ki67 staining showed markedly lower proliferation among cancer cells surrounded by CXCL16⁺ macrophages (Fig. 6d,e). Thus, CXCL16⁺ macrophages create a unique niche that tethers disseminated cancer cells and maintains transient quiescence.

**Fig. 6.**
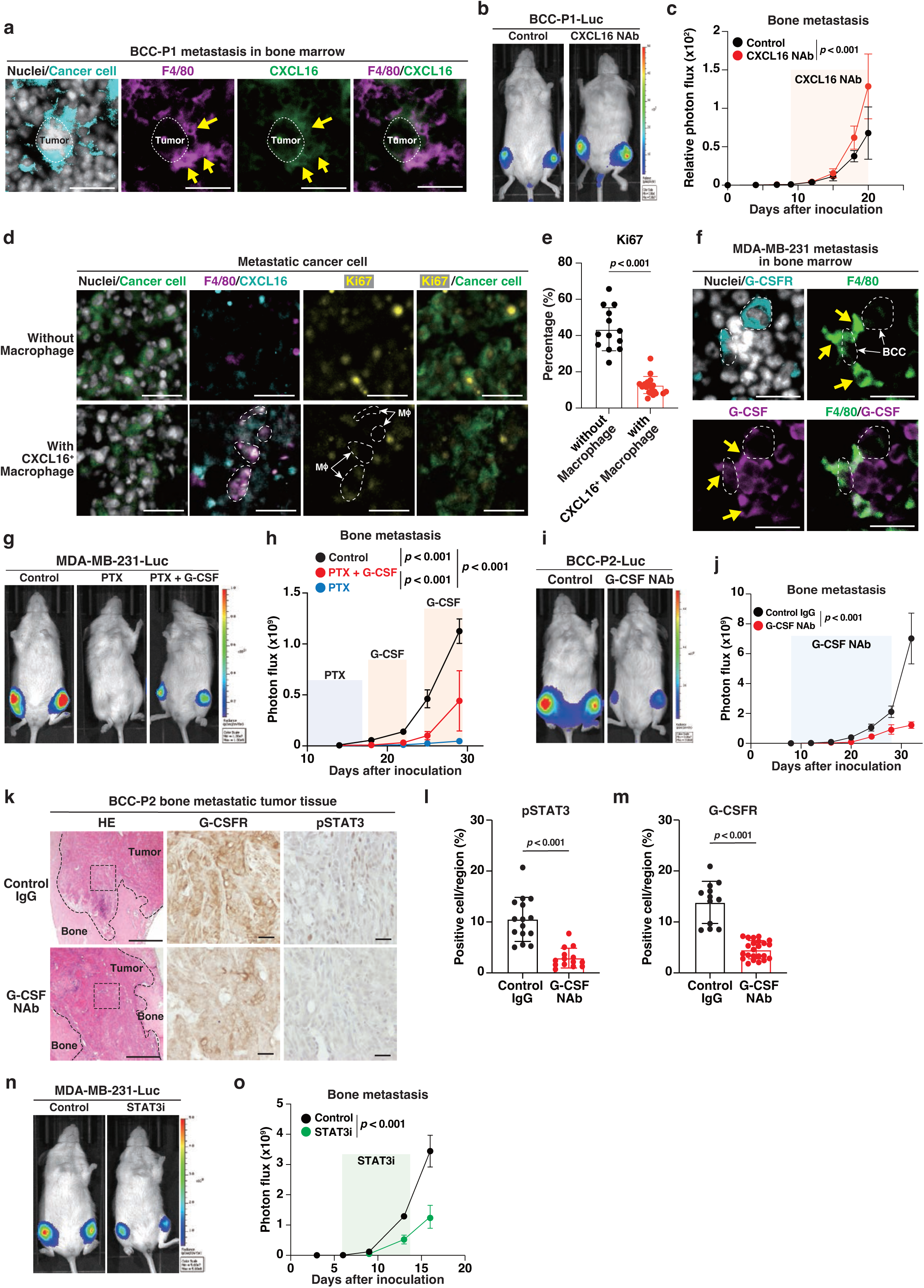
CXCL16^+^ macrophages niche imposes proliferative restraint in cancer cells and G-CSF^+^ macrophage niche supports metastasis-initiating cells. **a,** Representative images of bone metastatic tumors derived from BCC-P1 cells stained for cancer cells (STEM121, human-specific), F4/80 (macrophages), and CXCL16, with Hoechst counterstaining. Arrows indicate CXCL16⁺F4/80⁺ macrophages surrounding metastatic cancer cells. Scale bars, 100 µm. **b,** Representative IVIS images of bone metastases at Day 20. Beginning on Day 6, mice were treated with CXCL16-neutralizing antibody (CXCL16-NAb; 2.5 mg/kg) every other day for 20 days. **c**, Quantification of IVIS signals (mean ± SD; n = 3). **d**, Representative images of bone metastatic tumors derived from MDA-MB-231 cells stained for STEM121, F4/80, CXCL16, and Ki67; Hoechst counterstaining. Dotted lines and arrows indicate macrophages. Scale bars, 100 µm. **e**, Quantification of Ki67⁺ cancer cells in regions lacking macrophages versus regions containing CXCL16⁺ macrophages (mean ± SD; n = 13 and 19 random fields, respectively). **f**, Representative images of bone metastases stained for G-CSFR, F4/80, and G-CSF. Yellow arrows indicate G-CSF⁺F4/80⁺ macrophages adjacent to G-CSFR⁺ cancer cells (outlined by dotted lines; white arrows). Scale bars, 100 µm. **g,h**, IVIS images at Day 29 (**g**) and quantification (**h**) in mice treated with paclitaxel (PTX; 15 mg/kg intraperitoneally, three doses over 10 days) alone or followed by G-CSF (125 µg/kg intraperitoneally, daily for 5 days, two cycles). Data are mean ± SD (n = 3). **i,j**, IVIS images (**i**) and quantification (**j**) at Day 32 in mice treated with anti-G-CSF Nab (0.5 mg/kg intraperitoneally, once every two days). Data are mean ± SD (n = 3). **k**, H&E and immunohistochemistry for G-CSFR and pSTAT3. Scale bars, 500 µm (left) and 100 µm (right). **l,m**, Quantification of DAB-positive areas (mean ± SD; n = 3). **n, o**, IVIS images at day 16 (**n**) and corresponding quantification (**o**) of bone metastasis with or without STAT3 inhibitor (15 mg/kg, intraperitoneally, every other day for one week). Data are mean ± SD (n = 3 per group). Statistical significance: two-way ANOVA with Benjamini–Krieger–Yekutieli correction (**c, h, j, o**); unpaired two-tailed Student’s t-test (**e, l, m**).

In contrast, G-CSF⁺ F4/80⁺ macrophages appeared, adjacent to G-CSFR^high^ cancer cells in bone marrow lesions (Fig. 6f). G-CSF levels were elevated in metastasis-bearing bone marrow, and G-CSF expression correlated spatially with cancer-cell abundance (Extended Data Fig. 8c,d).

PTX suppressed bone metastasis when administered alone, but co-administration of G-CSF reinstated metastatic outgrowth (Fig. 6g,h). Clinical database analyses^70^ confirmed that high *G-CSFR* expression is associated with increased distant-metastasis risk (Extended Data Fig. 8e). Therapeutic disruption of the G-CSF–G-CSFR axis using either G-CSF NAb or the STAT3 inhibitor significantly reduced bone-metastatic burden and decreased G-CSFR and pSTAT3 levels in lesions (Fig.6i-o). Thus the G-CSF–G-CSFR axis promotes bone metastasis.

### CAFs orchestrate the metastatic niche comprising G-CSFR⁺ cancer cells and CXCL16⁺ macrophages

At late metastatic stages, CAFs accumulated abundantly within bone lesions in both PDX models and human clinical samples. G-CSFR^high^ cancer cells enriched at the borders of CAF-rich stroma, whereas CAF-poor tumor parenchyma contained few G-CSFR^high^ cells (Fig. 7a,d; Extended Data Fig. 9a; Supplementary Table 4). Quantitative analyses confirmed a significant positive correlation between α-SMA expression and G-CSFR fluorescence intensity (Fig. 7b,c,e,f).

**Fig. 7.**
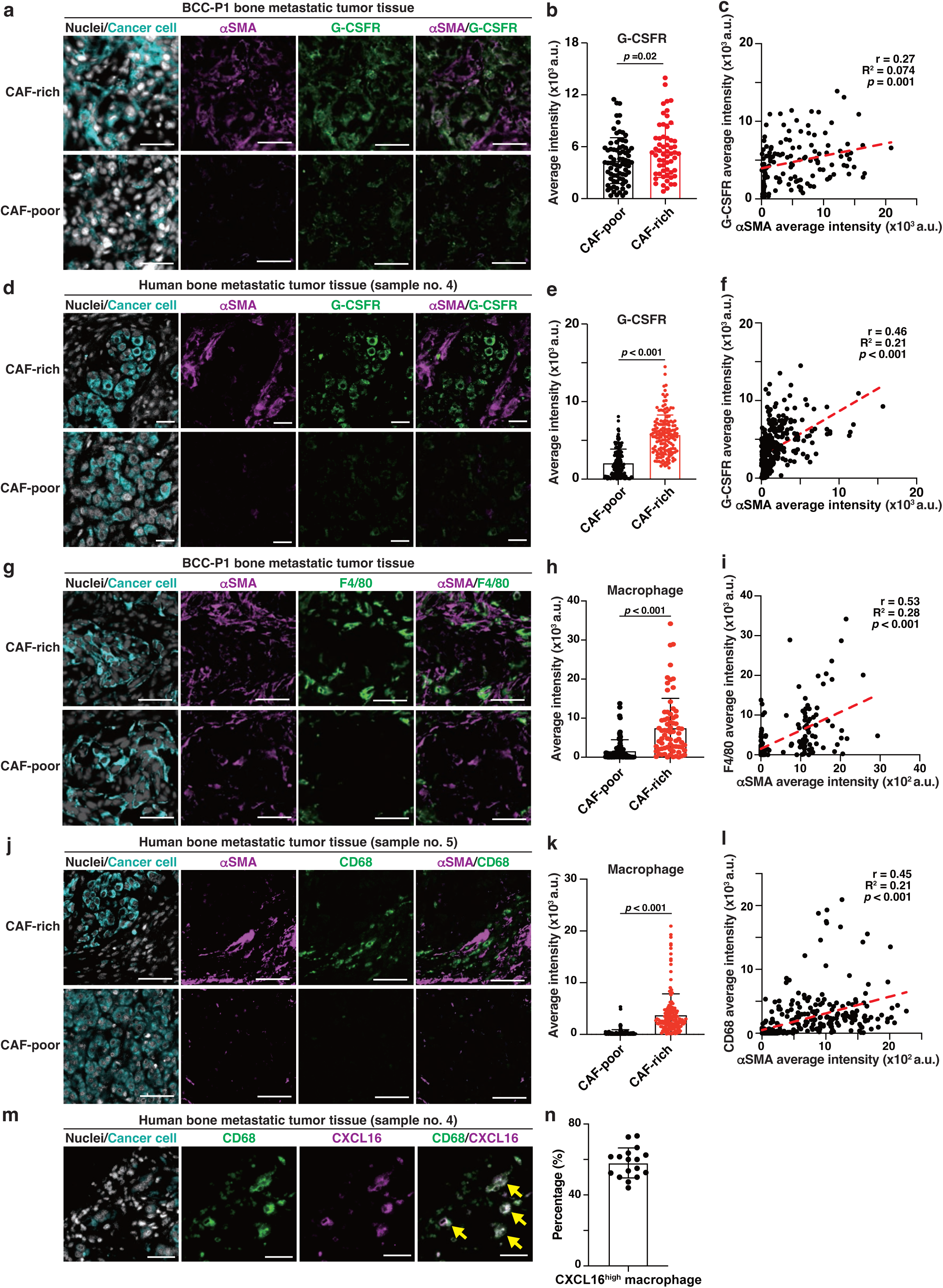
CAFs orchestrate a metastatic niche that nurtures G-CSFR^+^ metastasis initiating cells and CXCL16^+^ macrophages. **a, d,** Representative immunofluorescence images of bone metastatic tumors from PDX models (**a**) and human clinical samples (**d**), stained for cancer cells (STEM121; human specific), α-SMA, and G-CSFR. Nuclei were counterstained with Hoechst. Scale bars, 100 µm. **b, e,** Quantification of G-CSFR fluorescence intensity in CAF-poor versus CAF-rich regions. Data are mean ± SD (n = 77 CAF-poor and n = 58 CAF-rich fields in **b**; n = 93 CAF-poor and n = 84 CAF-rich fields in **e**). **c, f,** Pearson correlation between α-SMA and G-CSFR fluorescence intensities in bone metastases. **g, j,** Representative immunofluorescence images of bone metastatic tumors from PDX models (**g**) and human clinical samples (**j**), stained for cancer cells (STEM121), α-SMA, and macrophage markers F4/80 (mouse) in (**g**) and CD68 (human) in (**j**). Nuclei were counterstained with Hoechst. Scale bars, 100 µm. **h, k,** Quantification of macrophage fluorescence intensity—F4/80 (**h**) or CD68 (**k**) in CAF-poor and CAF-rich areas. Data are presented as mean ± SD (n = 83 CAF-poor and n = 71CAF-rich fields in **h**; n = 164 CAF-poor and n = 158 CAF-rich fields in **k**. **i, l,** Pearson correlation between α-SMA and macrophage (F4/80 or CD68) fluorescence intensities in bone metastases. **m,** Representative immunofluorescence images of human bone metastatic tumors stained for cancer cell (STEM121), CD68 (macrophages), and CXCL16. Nuclei were counterstained with Hoechst. Scale bars, 100 µm. **n,** Quantification of CXCL16⁺ macrophages among CD68⁺ cells. Data are presented as mean ± SD (n = 17 random fields). **b, e, h,** and **k,** Statistical significance was determined by unpaired two-tailed Student’s t-test.

By contrast, macrophages—identified by F4/80 in PDX models and CD68 in human samples—were restricted to the CAF-rich stromal compartment and absent from tumor parenchyma (Fig. 7g,j; Extended Data Fig. 9b), their signal intensities also correlated with α-SMA expression (Fig. 7h,i,k,l). Notably, among CD68⁺ macrophages in human bone metastases, approximately 60 % co-expressed CXCL16 (Fig. 7m,n).

Collectively, these data establish that CAFs orchestrate a composite metastatic niche that nurtures G-CSFR⁺ metastasis-initiating cancer cells and sustains CXCL16⁺ macrophages, integrating fibroblast- and myeloid-derived cues to support metastatic persistence and outgrowth.

## Discussion

This study reveals a previously unrecognized stromal–hematopoietic axis that shapes metastatic initiation, recurrence, and bone colonization in breast cancer. Using patient-derived systems, we show that tCAFs aberrantly secrete G-CSF, generating G-CSFR^high^ cancer cells primed for local relapse and skeletal dissemination (Fig. 8). Once in the bone marrow, DTCs cooperate with resident hematopoietic cells to assemble dynamic niches that determine whether metastatic growth is restrained or reactivated.

**Fig. 8.**
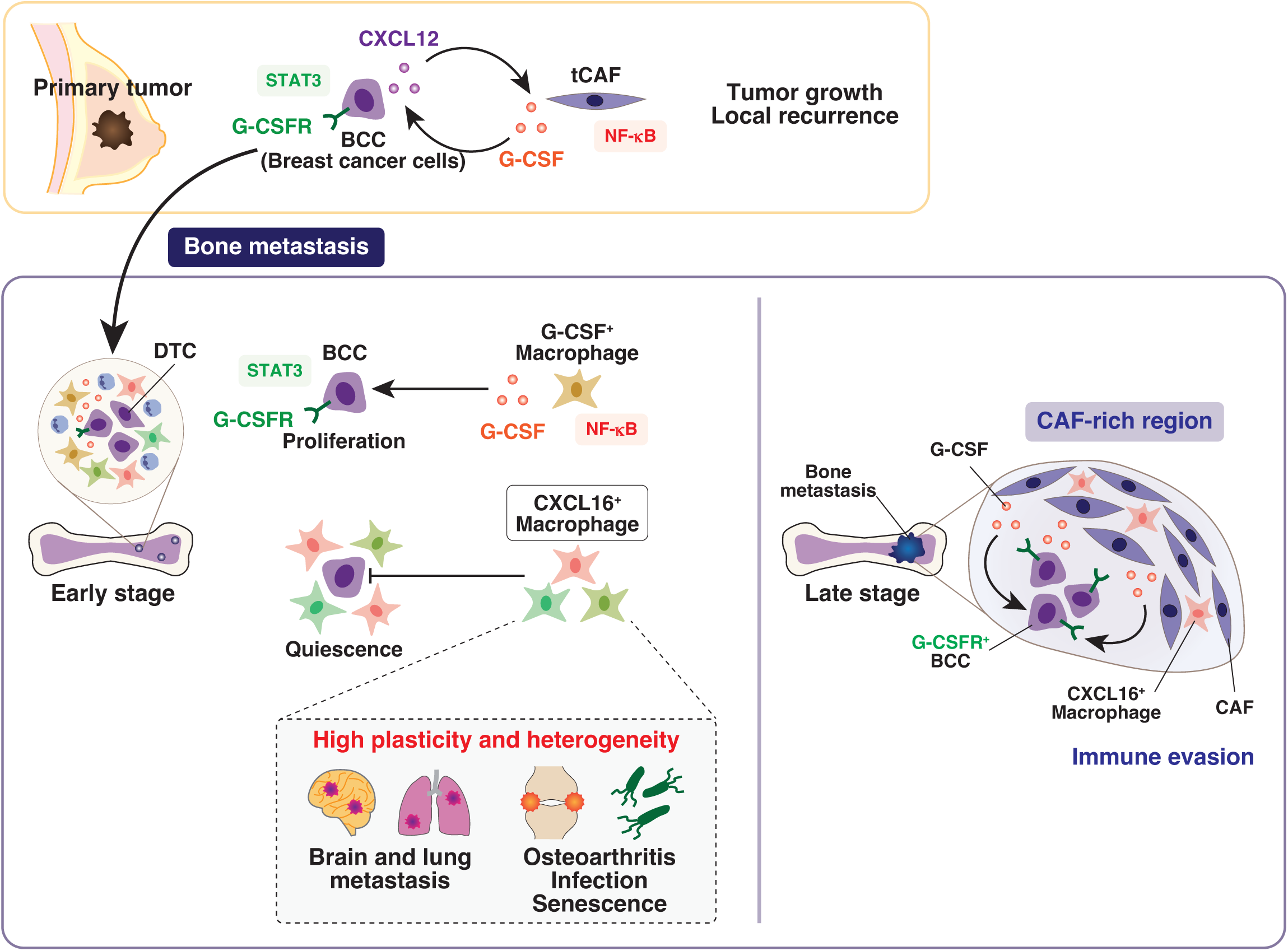
A stromal–hematopoietic circuit that fuels metastatic initiation and bone colonization. In primary tumors, NF-κB–activated tCAFs secrete G-CSF, reinforced by cancer-cell– derived CXCL12, generating G-CSFR^high^ cells primed for relapse. After entering bone marrow, DTCs encounter two opposing niches: a G-CSF⁺ macrophage niche promoting proliferation, and a CXCL16⁺ macrophage niche imposing transient restraint. CXCL16⁺ macrophages display marked plasticity, integrating TAM programs from distant metastases with non-tumor disease-associated macrophage signatures linked to infection, inflammation, autoimmune states, and senescence. In advanced lesions, they become confined to CAF-rich stroma, signaling a shift toward immune exclusion and evasion.

At the center of this network is a highly plastic CXCL16⁺ macrophage population—a transcriptionally heterogeneous and highly plastic state. Remarkably, the CXCL16⁺ macrophages integrate multiple TAM programs—including those characteristic of lung and brain metastases—alongside non-tumor disease–associated macrophage signatures related to infection, inflammation, autoimmune pathology, and cellular senescence. These CXCL16⁺ macrophages establish an early growth-restraining niche, whereas coexisting G-CSF⁺ macrophage states drive metastatic reactivation. Together with tCAF-derived G-CSF, these myeloid populations link fibroblast signaling to the temporal progression of metastasis. In advanced bone lesions, the confinement of CXCL16⁺ macrophages to CAF-rich stroma suggests a shift toward immune evasion.

Our single-cell analyses redefine the classical view of TAM polarization. CXCL16⁺ macrophages occupy a unique transitional state rather than fixed M1- or M2-like identities, integrating inflammatory and regulatory programs as well as non-tumor disease–associated signatures of acute and chronic inflammation. Notably, their transcriptional features resemble those of metastatic TAMs in distant organs, including lung and brain, suggesting that CXCL16⁺ macrophages may create “a metastasis-mimic niche” within the bone marrow. This extreme hybrid identity likely arises from local reprogramming of monocytes that respond to DTC invasion during the initial phase of bone metastasis, transiently restraining cancer-cell proliferation. The subsequent transition to G-CSF⁺ macrophages and neutrophils mark a shift from immune surveillance to a pro-metastatic state, underscoring macrophage plasticity as a key determinant of metastatic fate.

In advanced stages of bone metastasis in human samples, CXCL16⁺ macrophages were localized within CAF-rich stromal regions but were largely excluded from cancer-cell clusters. Consistent with previous reports showing that immune cells tend to be excluded from stromal regions of bone metastases^71^, this spatial segregation suggests that CAFs may recruit immune cells, including CXCL16⁺ macrophages, to the stromal compartment, thereby contributing to immune evasion. Given that CAFs aberrantly secrete cytokines such as G-CSF, these signals may not only attract myeloid cells but also reprogram them toward immunosuppressive or tumor-supportive phenotypes, reinforcing a stromal barrier that shields cancer cells from immune surveillance.

CXCL16 has dual roles as a membrane-bound adhesion molecule and a soluble chemokine with pleiotropic effects on immune cells^72^. Our data provide functional evidence that macrophage-derived CXCL16 contributes to transient tethering of disseminated cancer cells, facilitating their adaptation to the bone-marrow niche. Neutralization of CXCL16 abrogated this restraining effect, accelerating metastatic outgrowth. These findings indicate that early CXCL16⁺ macrophage niches act as temporary “landing pads” for DTCs, allowing phenotypic stabilization before the subsequent inflammatory switch. Given that metastasis-initiating cells must survive abrupt shifts in microenvironmental cues, such tethering interactions may represent a universal mechanism by which macrophages orchestrate the transition from dormancy to reactivation. Once DTCs lodge in the CXCL16⁺ niche, they may receive conflicting signals from coexisting TAM subsets, potentially endowing them with the capacity to adapt to other tissue microenvironments encountered during metastatic dissemination. Consistent with this hypothesis, breast-cancer patients with bone metastases often develop secondary distant metastases^3^, and prior studies indicate that cancer cells can be reprogrammed within bone marrow to acquire multi-organ metastatic competency^32^. Our future analyses will test this model directly.

In parallel, our data establish the CAF-derived G-CSF–G-CSFR axis as a molecular bridge linking primary and metastatic microenvironments. In primary tumors, tCAFs aberrantly secrete G-CSF in response to CXCL12 released from cancer cells, activating NF-κB signaling and reinforcing a paracrine loop that expands G-CSFR⁺ CSCs. These G-CSFR⁺ cells exhibit enhanced tumor-initiating capacity and pronounced chemoresistance and metastasis. In the bone marrow, macrophage- and neutrophil-derived G-CSF reactivates dormant metastasis-initiating cells, completing a fibroblast– myeloid circuit that underpins recurrence. Importantly, G-CSF is widely administered in oncology to prevent chemotherapy-induced neutropenia^26^. Our results raise caution that exogenous G-CSF may inadvertently expand G-CSFR⁺ cancer stem-lime cells, thereby increasing relapse risk—particularly in triple-negative breast cancer, where stromal G-CSF levels are intrinsically high. Indeed, several reports have linked tumor-cell–derived G-CSF expression to metastatic behavior; our findings extend this principle by showing that stromal G-CSF alone is sufficient to promote bone metastasis even when tumor-cell G-CSF levels are low.

Beyond its translational relevance, this study provides a conceptual framework for understanding how the bone marrow serves as a fertile soil for metastatic initiation. The emergence of G-CSF⁺ macrophage, CXCL16⁺ macrophage, and G-CSF⁺ neutrophil niches reveals that metastasis progresses through discrete microenvironmental phases—an early priming phase followed by a transient proliferative-restraint phase and, finally, an awakening phase—rather than a simple linear trajectory. CAF-rich stroma amplifies these transitions by sustaining both cancer stemness and myeloid reprogramming. The observation that CXCL16⁺ macrophages integrate TAM and non-tumor inflammatory programs suggests that the bone-metastatic niche co-opts tissue-repair and remodeling circuits normally engaged in homeostasis.

Collectively, our findings uncover an unappreciated layer of complexity in the interplay between fibroblasts, macrophages, and cancer cells. We propose that CAF–bone-marrow cross-talk acts as a central driver of metastatic initiation, coordinating cytokine and adhesion networks that dictate cancer-cell fate. Therapeutic strategies that selectively disrupt these stromal–myeloid interactions may prevent both local recurrence and distant relapse. Finally, our results highlight the need for judicious evaluation of clinical G-CSF use and open new avenues for targeting microenvironmental plasticity to combat therapy-resistant metastasis.

## Methods

### Human specimens

Fresh primary breast cancer tissues were obtained from patients undergoing surgical resection or biopsy at Kanazawa University Hospital and the University of Tokyo Hospital. The use of these samples was approved by the Institutional Review Boards of the Cancer Research Institute of Kanazawa University, Kanazawa University Hospital, the University of Tokyo Hospital (approval no. 72331-20).

Paraffin-embedded primary breast tumor tissues and bone metastatic tumor tissues were approved by the Institutional Review Board of Kanagawa Cancer Center (approval no. 28KEN4). Written informed consent was obtained from all patients prior to sample collection. All experiments were performed in accordance with the Declaration of Helsinki. All specimens were de-identified to protect patient confidentiality and were handled in compliance with national and international regulations.

### Establishment of patient-derived breast cancer cells (BCC-P1, P2, P3 and P4) and patient-derived cancer-associated fibroblasts (CAF-P5, P6, P7, P8 and P9)

To culture primary breast cancer cells (BCCs) and cancer-associated fibroblasts (CAFs), we treated the primary breast cancer tissues as described^16^. Primary breast cancer tissues were minced into ∼1-mm pieces and enzymatically dissociated with collagenase (Sigma-Aldrich, Merck, #C9407) and DNase I (100 U/mL; Sigma-Aldrich, Merck, #DN25) at 37 °C with gentle agitation. Cell suspensions were triturated, passed through 70-μm strainers, and treated with red blood cell lysis buffer (Roche, #11814389001).

To enrich tumor cells by magnetic negative selection, single-cell suspensions were incubated with a cocktail of biotin-conjugated antibodies consisting of a Lineage Cell Depletion Kit for hematopoietic/erythroid precursors (CD2, CD3, CD11b, CD14, CD15, CD16, CD19, CD56, CD123, and CD235a; Miltenyi Biotec, #130-092-211), anti-CD31 for endothelial cells (Thermo Fisher Scientific, #13-0310-82), and anti-PDGFRβ/CD140b for stromal/perivascular cells (BioLegend, #323604). After addition of anti-biotin microbeads, labeled cells were retained on MACS columns in a magnetic field, and the flow-through (Lin⁻/CD31⁻/PDGFRβ⁻) fraction was collected as the tumor-cell-enriched fraction according to the manufacturer’s instructions (Miltenyi Biotec).

Tumor-cell-enriched fractions were plated on collagen-coated dishes (IWAKI) and cultured either in organoid medium refer to the reported previously ^29,20^ or in EpiCult-C Human Medium Kit (STEMCELL Technologies, #05630) supplemented with the kit supplement mix, freshly prepared 1 μM hydrocortisone (Sigma-Aldrich, Merck, #H2270), 2 mM L-glutamine (Nacalai Tesque, #13004-02), 100 U/mL penicillin, and 100 μg/mL streptomycin (Nacalai Tesque, #26253-84). In parallel, the antibody-retained (CD31⁺ and/or PDGFRβ⁺) fraction-enriched for CAFs was eluted from the column and cultured in DMEM (Nacalai Tesque, #08458-45) supplemented with 20% fetal bovine serum (FBS; Thermo Fisher Scientific, #26140-079) and 1% penicillin–streptomycin (P/S).

### Cell culture and cell lines

Human breast cancer cell lines BT-20, BT-474, MDA-MB-231, and MCF7 were obtained from the American Type Culture Collection (ATCC, Manassas, VA, USA) and expanded upon receipt. BT-20 and BT-474 were maintained in RPMI-1640 (Nacalai Tesque, #30264-56) supplemented with 10% FBS and 1% penicillin/streptomycin (P/S). MDA-MB-231 and MCF7 were cultured in DMEM with 10% fetal bovine serum (FBS) and 1% P/S. mCherry-expressing CAFs (CAF-mCherry) were cultured in DMEM with 10% FBS and 1% P/S ^73^. Cells were maintained at 37 °C in a humidified atmosphere of 5% CO₂, and medium were replaced every 2–3 days. For passaging, cells were detached with Accumax (Innovative Cell Technologies, #AM105) or trypsin (Nacalai Tesque, #32777-44) for 3-10 minutes at 37 °C.

### Establishment of cell lines stably expressing HSV1-tk/eGFP/luc2

To monitor cancer cells in mice, BCC-P1, BCC-P2, and MDA-MB-231 cells were transduced with a triple-reporter construct expressing herpes simplex virus thymidine kinase 1 (HSV1-tk), enhanced GFP (eGFP), and firefly luciferase (luc2) genes, as previously described ^74^. The expression vector was kindly provided by Dr. Thordur Oskarsson (Department of Molecular Oncology, H. Lee Moffitt Cancer Center & Research Institute, Tampa, Florida, USA).

### Mice

Female NSG mice (NOD.Cg-Prkdc^scid Il2rg^tm1Wjl/SzJ; The Jackson Laboratory, stock no. 005557; RRID: IMSR_JAX:005557), 6–8 weeks old, were housed under specific pathogen– free conditions at 20–26 °C and 45–65% relative humidity on a 12-hour light/dark cycle, with ad libitum access to food and water. All animal procedures were approved by the Kanazawa University Animal Care and Use Committee (AP-24023) and performed in accordance with institutional and national guidelines.

Group sizes were chosen based on prior experience and pilot data; no statistical method was used to predetermine sample size. Animals were assigned to experimental groups prior to treatment using simple randomization to balance age and body weight. No animals were excluded from analyses unless prespecified humane endpoints were met; any exclusions are reported in the figure legends.

### Preparation of conditioned medium (CM)

To prepare CM derived from CAFs cultured alone, cells were seeded and, 12 hours later, the medium was replaced with serum-free medium. After 24 hours, the supernatant was collected and passed through a 0.45-μm filter (Millipore Sigma, Merck) to remove cell debris.

To prepare CM derived from CAFs co-cultured with BCCs, CAFs were seeded in a culture dish and separately BCCs were seeded onto Transwell inserts (Corning). At 12 hours after seeding, the BCCs-culture Transwell inserts were placed into the CAFs-culture dishes to initiate co-culture in the serum-free medium, and incubation continued for 72 hours. CM was then collected and filtered through a 0.45-μm filter.

### Tumor spheroid formation assay

Breast cancer cells were cultured under non-adherent, serum-free conditions to assess growth in a sphere assay. Briefly, single-cell suspensions were seeded into ultra–low-attachment plates (Corning) in the sphere culture medium^16^. CM derived from CAF-P5 co-cultured with BCC-P1 was added to the cultures, with or without neutralizing anti-G-CSF antibody (5 µg/µL; R&D Systems, #AF-214-NA). BCC cells were cultured in the presence of human recombinant G-CSF (0.5 or 1.0 µg/mL; Thermo Fisher Scientific, #AF-300-23) for 7 days under spheroid conditions. Spheroid formation was monitored, and bright-field images were acquired using an inverted microscope (Nikon). Sphere number and size were quantified with ImageJ software (NIH).

Spheroid-forming ability was evaluated by extreme limiting dilution assays (ELDA). Cells were plated at serial dilutions into ultra–low-attachment 96-well plates in sphere culture medium, and wells containing spheres with a diameter ≥75 µm were scored as positive after the culture period. Stem cell frequency and statistical significance were calculated using the ELDA software (Walter and Eliza Hall Institute; http://bioinf.wehi.edu.au/software/elda/).

### RNA-sequencing analysis

Bulk RNA-seq was performed on CAFs cultured either alone or co-culture with BCC-P1, using three independent CAF lines (CAF-P5, CAF-P8, and CAF-P9). Bulk RNA-seq was also performed on breast cancer cells cultured either alone or co-culture with patient-derived CAFs, using three independent BCC lines (BCC-P1, BCC-P3, and BCC-P4).

Total RNA was extracted using the NucleoSpin RNA XS kit (Macherey-Nagel) according to the manufacturer’s instructions. cDNA synthesis and amplification were performed with the SMARTer Ultra Low Input RNA Kit for Sequencing (Takara Bio) using ≤ 10 ng total RNA and no more than 12 PCR cycles to minimize amplification bias. cDNA quality was assessed on an Agilent 2100 Bioanalyzer using the High Sensitivity DNA Kit (Agilent Technologies). Illumina libraries were prepared and sequenced to generate ∼90 million paired-end reads (2 × 101 bp) per library on a HiSeq 2000/2500 (Illumina).

Adapters and low-quality bases were removed prior to alignment. Trimmed reads were mapped to the human reference genome hg19 (UCSC, build 20150519) with TopHat2 (v2.0.10), and transcripts were assembled with Cufflinks (v2.1.1) using RefSeq annotations. Gene expression was quantified as fragments per kilobase of transcript per million mapped reads (FPKM) with Cuffdiff (v2.1.1). Unless otherwise stated, differentially expressed genes were defined at false discovery rate (FDR) q < 0.05 (Benjamini–Hochberg). Differential expression results were visualized as volcano plots (log₂ fold change versus −log₁₀ adjusted P value).

To evaluate pathway-level differences, gene set enrichment analysis (GSEA) was performed specifically for the comparison between CAFs co-cultured with BCCs and those cultures alone. Ranked gene lists were generated from bulk RNA-seq differential expression results, ordered by log₂ fold change between CAFs co-cultured with BCCs and those cultures alone. Analyses were conducted using the GSEA desktop application (Broad Institute, v4.3.2) with gene sets obtained from the Molecular Signatures Database (MSigDB, v7.5.1). The Hallmark (H) curated (C2), and immunologic (C7) gene set collections were used as reference databases. Statistical significance was determined based on 1,000 phenotype-based permutations. Gene sets were considered significantly enriched when the FDR q-value was less than 0.25, in accordance with the GSEA default criteria. For selected analyses, a more stringent cutoff of FDR q < 0.05 was applied, as indicated in the figure legends. Normalized enrichment score (NES), nominal P value, and FDR q-value were reported for each enriched pathway. Enrichment plots, leading-edge subsets, and summary reports were generated by the GSEA desktop software, and representative figures were exported for visualization.

### Quantitative real-time PCR (qPCR)

Total RNA was extracted from cells or tissue samples using QIAshredder and the RNeasy Mini Kit (Qiagen) according to the manufacturer’s instructions. RNA purity and concentration were measured with a NanoDrop 2000 spectrophotometer (Thermo Fisher Scientific). cDNA was synthesized using the High-Capacity cDNA Reverse Transcription Kit (Thermo Fisher Scientific). Primers were designed with Primer-BLAST (NCBI) and validated for specificity and efficiency. The following primer pairs were used: *CSF3-1* (forward: 5’-CCAGGAGAAGCTGGTGAGTGAG-3’, reverse: 5’-CGGGGTGGCACAGCTTGTAG-3’), *CSF3-2* (forward: 5’-AGGAGAAGCTGGTGAGTGAGT-3’, reverse: 5’-AGGGGATGCCCAGAGAGTGTC-3’), *CXCL12-1*(forward: 5’-GCTACAGATGCCCATGCCGA-3’, reverse: 5’-AGTCAGAGGAGTGGCTCCGT-3’), *CXCL12-2* (forward: 5’-ATGAACGCCAAGGTCGTGGT-3’, reverse: 5’-TCGGCATGGGCATCTGTAGC-3’), *CFS3R* (forward: 5’-GCTCAAGATCACAAAGCTGGT-3’, reverse: 5’-CCGCACTCCTCCAGACTTC-3’), *ACTB* (forward: 5’-AAGTCCCTTGCCATCCTAAAA-3’, reverse: 5’-ATGCTATCACCTCCCCTGTG-3’) and *GAPDH*(forward: 5’-AACTTTGGCATTGTGGAAGG-3’, reverse: 5’-ACACATTGGGGGTAGGAACA-3’) as the reference gene. Quantitative PCR was performed using Fast SYBR Green Master Mix (Thermo Fisher Scientific) on an Applied Biosystems QuantStudio 1 Real-Time PCR System. Gene expression was calculated by the comparative Ct method (2^−ΔΔCt), normalized to *ACTB* or *GAPDH* as endogenous controls. All reactions were run in technical triplicates, and data represent at least three independent experiments.

### Enzyme-linked immunosorbent assay (ELISA)

Concentrations of G-CSF in CM were measured by Quantikine ELISA kit (Human G-CSF; R&D Systems) according to the manufacturer’s instructions. Briefly, 100 µL of diluted CM was added per well to 96-well plates pre-coated with captured antibodies and incubated for 2 hours at room temperature. Plates were washed with the kit wash buffer, 100 µL per well of horseradish peroxidase–conjugated detection antibody was added, and plates were incubated for 1 hour at room temperature. After four washes, 100 µL of tetramethylbenzidine (TMB) substrate solution was added and the reaction was developed for 20 minutes in the dark, then stopped with 50 µL of 2N sulfuric acid. Absorbance was read at 450 nm with wavelength correction at 570 nm using a SpectraMax iD3 microplate reader (Molecular Devices). Standard curves were generated using Human G-CSF standards provided in the kit, and concentrations were calculated by interpolation. All samples were measured in duplicate, and experiments were repeated three times independently.

### Western blotting

Western blotting was performed as described previously^75^. Briefly, cells were washed with Phosphate buffered saline (PBS) and lysed on ice for 5 minutes in RIPA buffer (Thermo Fisher Scientific, #89901) supplemented with 0.5 mM EDTA, 1 mM DTT, 0.5 mM PMSF, protease inhibitor cocktail (Nacalai Tesque, #25955-11), and phosphatase inhibitors (Nacalai Tesque, #07575-51). Lysates were rotated at 4 °C for 60 minutes and clarified by centrifugation at 15,000 × g for 10 minutes at 4 °C. Protein concentrations were determined using the Bradford Assay (Bio-RAD) according to the manufacturer’s instructions. Equal amounts of protein samples were mixed with Laemmli sample buffer and heated at 95°C for 5 minutes, resolved by sodium dodecyl sulfate-polyacrylamide gel electrophoresis (SDS-PAGE) on 10% polyacrylamide gels, and transferred to PVDF membranes (Merck Millipore). Membranes were activated in methanol, rinsed, and blocked for 1 hour at room temperature in PBST (PBS + 0.1% Tween-20) containing 5% skim milk.

Blocked membranes were incubated overnight at 4 °C with primary antibodies diluted in 5% skim milk in PBST. The following antibodies were used: anti-α-SMA (1:2000; Abcam, #ab5694), E-Cadherin (1:1000; Cell Signaling Technology, #3195), anti-N-Cadherin (1:2000; BD Biosciences, #610920), anti-Vimentin (1:1000; Cell Signaling Technology, #5741), and anti-β-actin (1:5000; Sigma-Aldrich, Merck, #MAB1501) as a loading control. After three washes in PBST, membranes were incubated with horseradish peroxidase (HRP)-conjugated secondary antibodies (anti-rabbit IgG, 1:5000, Cytiva, #NA934; anti-mouse IgG, 1:5000, Cytiva, #NA9310) for 1 hour at room temperature. Signals were developed with Enhanced Chemiluminescence (ECL) Prime Western Blotting Detection Reagent (Sigma-Aldrich, Merck) on a ChemiDoc Imaging System (Vilber). Band intensities were quantified using ImageJ software (NIH), and protein expression levels were normalized to β-actin.

### CAF subtype deconvolution and *G-CSF* expression analysis using publicly available scRNA-seq data set

CAF subtype composition was inferred from bulk RNA-seq using MuSiC (R package, v3.0)^76^. As a single-cell reference, we used the publicly available dataset^77^, restricting the reference to CAF clusters and their subtype annotations as defined in the original study. Cells with low complexity or high mitochondrial RNA content were removed, and gene expression matrices were normalized as in the reference publication. For each donor in the reference, gene-wise averages were computed per CAF subtype to enable subject-aware weighting.

For deconvolution, the bulk RNA-seq input consisted of CAFs in CAF-P5, CAF-P8, and CAF-P9 co-cultured with BCC-P1. To mitigate library-size effects, bulk expression values were converted from FPKM to TPM prior to deconvolution (TPM_i = FPKM_i / ΣFPKM × 10^6^). Only genes shared between the bulk and reference datasets were retained; genes with low expression (e.g., bulk CPM < 0.1) or expressed in < 10% of cells within a reference subtype were excluded. Cell deconvolution was performed using *music_prop* function in MuSiC with default parameters.

Additionally, using the same single-cell reference dataset, we quantified expression of *CSF3* (*G-CSF*) across CAF subtypes. Results were visualized as dot plots.

### Immunocytochemistry

Immunocytochemistry was performed as described previously^75^. Briefly, CAFs co-cultured with BCCs and those cultured alone were seeded on collagen-coated 8-well chamber glass slides (Corning). 24 hours later, cells were fixed with 4% paraformaldehyde (PFA) (Nacalai Tesque, #09154-85) in PBS for 15 minutes, permeabilized in 0.5% Triton X-100 (Nacalai Tesque, #25987-85) in PBS for 15 minutes, and blocked with 1% BSA (Nacalai Tesque, #01863-48) in PBS for 1 hour at room temperature. Then cells were incubated with the primary antibodies at optimized concentrations at 4 °C overnight. Primary and secondary antibodies were diluted in 1% BSA/PBS: anti-NF-κB (1:200; Cell Signaling Technology, #4764), and anti-G-CSFR (1:200; Novus, #NBP2-61696). After three washes in PBS, cells were incubated with secondary antibodies (Alexa Fluor 488 goat anti-rabbit, 1:200, Thermo Fisher Scientific, #A-11008; Alexa Fluor 488 goat anti-mouse, 1:200, Thermo Fisher Scientific, #A-11001) for 1 hour at room temperature in the dark. Nuclei were stained with Hoechst (1:2500; Thermo Fisher Scientific, #H3570) for 30 minutes. Slides were mounted with fluorescence mounting medium (Agilent, #S3023). Images were acquired by using Zeiss LSM900 confocal microscopy and were analyzed by ImageJ software.

### Mix organoid culture and immunostaining

BCC-P1 cells were dissociated with Accumax (Innovative Cell Technologies) for 5-10 minutes at 37 °C. CAF-mCherry cells were dissociated with trypsin (Nacalai Tesque) for 3 minutes at 37 °C. Single cells were collected and resuspended in phosphate-buffered saline (PBS; Nacalai Tesque).

Cell suspensions were prepared at a total density of 2 × 10^4^ cells, mixed at a ratio of BCCs to CAFs of 2:1, and combined with 1.2% methylcellulose and organoid culture medium^29,20^ to a final volume of 500 µL. 20 µL droplets of this mixture were plated in an inverted orientation and incubated overnight at 37 °C. The formation of cellular aggregates within droplets was confirmed, and each aggregate was subsequently embedded in 50-70 µL Matrigel. After solidification for 20 minutes at 37 °C, organoid medium was added.

Immunostaining was performed following the protocol described previously^78^. Briefly, organoids were incubated for 10 minutes at 4 °C with Cell Recovery Solution (Corning, #354253) and fixed in 4% PFA in PBS for 30 minutes at room temperature. Samples were permeabilized with solution A (40% CUBIC-L [Tokyo Chemical Industry, #T3740], 5 mM NaCl, and 10 µg/mL DAPI [Dojindo, #340-07971] in distilled water) at 37 °C overnight, followed by incubation with 50% CUBIC-R (Tokyo Chemical Industry, #T3983) in distilled water for 15 minutes and then incubated in 100% CUBIC-R for 15 minutes at room temperature.

Samples were incubated for 5 days at 4 °C with primary antibodies, including anti-G-CSFR (1:50, Novus), diluted in 1 M HEPES-TSC (pH 7.5) buffer. After washing, samples were incubated with species-appropriate Alexa Fluor-conjugated secondary antibodies (1:50, Thermo Fisher Scientific) for 2 days at 4 °C. Organoids were washed with 0.1 M PBT and 0.1 M PB before mounting in Mounting Solution (Tokyo Chemical Industry, #M3294). Images were acquired using Zeiss LSM900 confocal microscopy for all conditions and analyzed with ImageJ software.

### Primary breast cancer xenograft mouse model

For orthotopic xenotransplantation, patient-derived BCCs (BCC-P1: 3 × 10^5^ cells; BCC-P2-Luc: 2 × 10^5^ cells) suspended in 100 µL of a 1:1 mixture of PBS and Matrigel (Corning) were injected into the fourth mammary fat pads of female NSG mice. When tumors reached approximately 500 mm³, they were surgically resected. We determined the doses of G-CSF NAb, PTX, G-CSF and STAT3 inhibitor based on the literature (G-CSF NAb^79^, G-CSF^80^, PTX^20^, STAT3 inhibitor^81^.

For G-CSF neutralizing antibody (G-CSF NAb) treatment, mice received intraperitoneal injections of G-CSF NAb (0.5 mg/kg; R&D Systems, #MAB414) once every 2 days. Control groups received intraperitoneal injections of control IgG antibody (0.5 mg/kg; R&D Systems, #MAB005) on the same schedule.

For paclitaxel (PTX) and/or G-CSF treatment, mice were divided into three groups: control, PTX alone, and PTX followed by G-CSF. PTX (10 mg/kg; FUJIFILM Wako, #167-28166) was administered intraperitoneally twice per week for 2 weeks after tumor formation was confirmed. G-CSF (125 µg/kg; Thermo Fisher Scientific, #AF-300-23) was then administered intraperitoneally once daily for 5 days, for two consecutive cycles following PTX treatment. When control tumors reached approximately 2,000 mm³ or tumors of treatment groups were monitored until 140 days, they were surgically resected. For the treatment of recurrent tumor, the STAT3 inhibitor C188-9 (100 mg/kg; Selleck, #S8605) was administered intraperitoneally once daily for 1 week, for two consecutive cycles.

Primary tumor growth was monitored using a digital caliper and/or bioluminescence imaging. Tumor volume was calculated using the following formula: V = L × L × S / 2, where L and S represent the lengths of the major and minor axes, respectively. Body weight was monitored every 2–3 days. For tumor monitoring by bioluminescence imaging, mice were anesthetized with 2% isoflurane (Viatris, #871119) in oxygen and injected intraperitoneally with D-luciferin (150 mg/kg; FUJIFILM Wako, #126-05116). Images were captured using an IVIS Spectrum system (IVIS Lumina LT Series III, PerkinElmer). Bioluminescence was analyzed using Living Image software v.4.4 (Caliper Life Sciences).

### Bone metastatic xenograft model

To create bone metastasis model, patient-derived BCCs expressing luciferase (BCC-P1: 3 × 10^5^ cells; BCC-P2-Luc: 5 × 10^5^ cells) or MDA-MB-231-Luc cells (5 × 10^5^ cells) suspended in 100 µL PBS were injected into the caudal (tail) artery of female NSG mice^82^.

For G-CSF neutralizing antibody (G-CSF NAb) treatment, mice received intraperitoneal injections of G-CSF NAb (0.5 mg/kg; R&D Systems) once every 2 days. Control groups received intraperitoneal injections of control IgG antibody (0.5 mg/kg; R&D Systems) on the same schedule.

For PTX and/or G-CSF treatment, mice were divided into three groups: control, PTX alone, and PTX followed by G-CSF. PTX (15 mg/kg; FUJIFILM Wako) was administered intraperitoneally twice per week for 2 weeks after tumor formation was confirmed. G-CSF (125 µg/kg; Thermo Fisher Scientific) was then administered intraperitoneally once daily for 5 days, for two consecutive cycles following PTX treatment. For the treatment of metastatic tumor, the STAT3 inhibitor C188-9 (50 mg/kg; Selleck) was administered intraperitoneally every other day for 1 week.

For CXCL16 neutralizing antibody (CXCL16 NAb) treatment, we determined the doses of CXCL16 NAb based on the literature^83^. Mice were received intraperitoneal injections of CXCL16 NAb (2.5 mg/kg; R&D Systems, #MAB503) once every 2 days. Control groups received intraperitoneal injections of control IgG antibody (2.5 mg/kg; R&D Systems, #MAB006) on the same schedule.

At the experimental endpoint, mice were euthanized, and femurs were excised. Bones were fixed in 10% neutral buffered formalin (Nacalai Tesque, #37152-51) for 24 hours and processed for paraffin embedding or fixed in 4% paraformaldehyde (PFA, Nacalai Tesque) in PBS and embedded in OCT compound (SAKURA, #4583) for frozen sectioning. Bones were decalcified in 0.5 M EDTA (pH 7.4) followed by sequential incubation in a discontinuous sucrose gradient (5%, 10%, 15%, and 20%) for 12 hours each prior to embedding^84^.

### Hematoxylin and eosin (HE) staining

Paraffin-embedded sections (4 µm) were deparaffinized by immersion in xylene and rehydrated through a descending ethanol series to distilled water. Nuclei were stained with Mayer’s hematoxylin solution (FUJIFILM Wako, #131-09665) for 9 minutes. Cytoplasmic staining was performed with 1% eosin Y solution (FUJIFILM Wako, #051-06515) for 2 minutes. Sections were dehydrated through an ascending ethanol series, cleared in xylene and mounted with Multi-Mount 48 (Matsunami, #FM48001).

### Immunohistochemistry

#### Paraffin-embedded sections and DAB staining

Immunohistochemistry was performed as described previously^75^. Briefly, resected tumor tissues were fixed in 10% neutral buffered formalin (Nacalai Tesque) and embedded in paraffin. Sections (5-10 µm) were deparaffinized, rehydrated, and subjected to antigen retrieval using citrate buffer (Target Retrieval Solution, Agilent, pH 6.0 [#S2369] for GCSFR; pH 9.0 [#S2367] for phospho-STAT3) at 121 °C for 10 minutes. Endogenous peroxidase activity was quenched with 3% hydrogen peroxide in methanol for 15 minutes. After blocking with Protein Block Serum-Free (Agilent, #X0909) for 1 hour, sections were incubated with primary antibodies overnight at 4 °C. The primary antibodies used in this experiment are as follows: anti-G-CSFR antibody (1:200; Novus, #NBP2-61696), anti–phospho-STAT3 antibody (1:50; Cell Signaling Technology, #9145). Detection was performed using HRP-linked secondary antibody reagents (Histofine Mouse MAX-PO (R) or Rat MAX-PO (M), Nichirei Bioscience Inc.) and visualized with DAB chromogen (Nichirei Bioscience Inc.). Slides were counterstained with hematoxylin (FUJIFILM Wako), dehydrated through graded ethanol, cleared in xylene (Nacalai Tesque, #36611-03), and mounted with Multi-Mount 480 (Matsunami). Images were captured using an Olympus IX83 inverted microscope. Positive staining was quantified in 5-7 randomly selected fields per section using ImageJ software.

#### Immunofluorescent staining of frozen tumor tissue

Immunohistochemistry was performed as described previously^75^. Briefly, tumor tissues were fixed in 4% PFA (Nacalai Tesque) in PBS overnight and cryoprotected by sequential incubation in 30%, 20%, and 10% sucrose solutions (FUJIFILM Wako, #196-00015) at 4 °C for 12 hours each. Samples were embedded in Tissue-Tek OCT compound (SAKURA) and cut into 8-12 µm cryosections using a Leica CM1950 cryostat.

Frozen tumor tissue sections were fixed in pre-cooled acetone (−20 °C; FUJIFILM Wako, #016-00346) until evaporation, followed by melting of the OCT compound at room temperature. Sections were permeabilized with 0.5% Triton X-100 (Nacalai Tesque) in PBS for 30 minutes at room temperature, then blocked with 1% BSA (Nacalai Tesque) in PBS for 1 hour at room temperature. Sections were incubated with primary antibodies at optimized concentrations overnight at 4 °C, followed by incubation with fluorophore-conjugated secondary antibodies at appropriate dilutions for 1 hour at room temperature. Nuclei were counterstained with Hoechst 33342 (Thermo Fisher Scientific).

#### Immunofluorescent staining of frozen bone tissue

Frozen bone tissue sections were air-dried for 10 minutes at room temperature to remove OCT. Sections were blocked with 10 % BSA (Nacalai Tesque) in PBS for 1 hour at room temperature, followed by incubation with primary antibodies at overnight at 4 °C. After washing, sections were incubated with fluorophore-conjugated secondary antibodies for 1 hour at room temperature. Nuclei were counterstained with Hoechst 33342 (Thermo Fisher Scientific).

#### Immunofluorescent staining of formalin-fixed, paraffin-embedded (FFPE) human clinical tumor tissues

FFPE clinical tumor specimens (5 μm) were deparaffinized, rehydrated, and subjected to heat-induced epitope retrieval (pH 6.0 for GCSFR; pH 9.0 for CD68, CXCL16) in a pressure cooker. Endogenous peroxidase was blocked with Protein Block Serum-Free (Agilent) for 1 hour, sections were incubated overnight at 4 °C with primary antibodies. Washed with PBS three times, and then incubated with species-appropriate, highly cross-adsorbed fluorophore-conjugated secondary antibodies (1 hour, room temperature, protected from light). Nuclei were counterstained with Hoechst 33342 (Thermo Fisher Scientific), and slides were mounted in Fluorescence Mounting Medium (Agilent) for fluorescence microscopy. Images were captured using a confocal microscope (Keyence BZ-X800) under identical settings for all conditions and analyzed using ImageJ.

#### Antibodies

The following primary antibodies were used: STEM121 (1:100; Takara Bio, #Y40410), anti-G-CSFR (1:200; Novus, #NBP2-61696), anti-α-SMA (1:100; Abcam, #ab5694), anti-G-CSF (1:100; Abcam, #ab181053), anti-mouse CXCL16 (1:100; R&D Systems, #AF503), and anti-mouse F4/80 (1:100; BioLegend, #123102), anti-human CD68 (1:100; Thermo Fisher Scientific, #14-0681-82), anti-human CXCL16 (1:20; R&D Systems, #AF976), anti-Ki-67 (1:400; Cell Signaling Technology, #9129), anti-CD31 (1:100; Thermo Fisher Scientific, #13-0310-82). To achieve direct labeling, the Zenon Mouse IgG Labeling Kit (Alexa Fluor 647, Thermo Fisher Scientific, #Z25008) was used with STEM121 antibody (final concentration 1:100), and the Zenon Rabbit IgG Labeling Kit (Alexa Fluor 647, Thermo Fisher Scientific, #Z25308) was used with anti–α-SMA antibody (final concentration 1:100), according to the manufacturers’ instructions. Images were acquired using a KEYENCE BZ-X800 fluorescence microscope under identical settings for all conditions and analyzed with ImageJ software (NIH).

### Flow cytometry and cell sorting

BCCs were dissociated into single-cell suspensions and resuspended in fluorescence activated cell sorting (FACS) buffer (PBS supplemented with 0.5% BSA and 1 mM EDTA) at a concentration of 5 × 10^6^ cells/ml. Cells were stained with anti-human G-CSFR antibody (1:80; Novus), followed by a PE-Cy7-conjugated secondary antibody (1:80; Thermo Fisher Scientific, #12-4010-82) for 30 minutes at 4 °C in the dark. After staining, cells were washed twice with FACS buffer and incubated with 7-aminoactinomycin D (7-AAD; 1:5, BD Biosciences, #555815) for 10 minutes at 4 °C to exclude dead cells.

Cell sorting was performed on a FACSAria III cell sorter (BD Biosciences) equipped with a 100-µm nozzle. Based on G-CSFR fluorescence intensity, the top 15% (G-CSFR^high^) and bottom 15% (G-CSFR^low^) populations were isolated for subsequent assay.

### Drug treatment using organoid culture

BCC-P1 cells were dissociated with Accumax (Innovative Cell Technologies) for 5-10 minutes at 37 °C. Single cells were collected and resuspended in organoid medium. A total of 8 × 10^5^ cells were embedded in 150 µL Matrigel. After solidification for 30 minutes at 37 °C, organoid medium was added. Organoid growth was monitored using an inverted phase-contrast microscope (Nikon ELWD 0.3 T1-SNCP).

For drug treatment, organoids were treated with vehicle (0.1% DMSO) or Paclitaxel (PTX) (10 µM, FUJIFILM Wako) for 5 days as a positive control. G-CSF (10 ng/mL, Thermo Fisher Scientific) were added to the culture medium on day 3 together with PTX, and organoids were further cultured for 3 days. For endpoint analysis, organoids were dissociated with TrypLE Express (Thermo Fisher Scientific, #12605-010) and trypsin (Nacalai Tesque), and single cells were resuspended in FACS buffer.

Cells were first stained with anti-human G-CSFR antibody (1:200; Novus) followed by PE-conjugated secondary antibody (1:80; Thermo Fisher Scientific) for 30 minutes at 4°C in the dark. After two washes, cells were resuspended in FACS buffer with 7-Aminoactinomycin D (7-AAD, 1:5; BD Biosciences) for 10 minutes to exclude dead cells. Cells were then fixed and permeabilized using the Foxp3/Transcription Factor Staining Buffer Set (Thermo Fisher Scientific), according to the manufacturer’s instructions to enable intracellular staining for Ki67. Cells were stained with anti-human Ki67 antibody (1:20; Thermo Fisher Scientific, #1404-5698-82) conjugated to Brilliant Violet 421 for 30 minutes at room temperature in the dark. Flow cytometry was performed on a SONY Cell Sorter MA900. Data were analyzed using FlowJo software (v10.10.0). A minimum of 10,000 events acquired per sample. The percentages of G-CSFR-positive and Ki67-positive cells were quantified, and co-expression was assessed by quadrant analysis. Experiments were performed in triplicate.

### Kaplan–Meier survival analysis

Kaplan–Meier survival analysis was conducted using the PrognoScan database (http://dna00.bio.kyutech.ac.jp/PrognoScan/). Breast cancer datasets GSE1379 and GSE7390 were selected for relapse-free survival (RFS) and distant metastasis-free survival (DMFS) analyses. Patients were dichotomized into high- and low-expression groups based on the median expression level of G-CSFR (CSF3R), and the medians were used as the cutoff values. Survival curves were generated using the Kaplan-Meier method in GraphPad Prism (v10.5.0), and statistical significance between groups was assessed by the log-rank (Mantel-Cox) test.

### Single-cell RNA sequencing (scRNA-seq)

Bone samples were collected at day 10 and day 20 post-injection of 3 × 10^5^ patient-derived breast cancer cells (BCC-P1) via tail artery injection of NSG mice (n = 2 mice for control and n = 3 mice each for Day 10 and Day 20).

#### Bone marrow cell isolation

Bone marrow cells were isolated as described^84^. Briefly, femurs were aseptically dissected, and each femur was flushed using 1 mL of Accumax (Innovative Cell Technologies) and 0.1 mg/mL DNase I (Merck) with a 21-gauge needle. The suspension was gently inverted to confirm tissue dispersion, followed by incubation with rotation at 26 °C for up to 20 minutes using a MACSmix rotator (Miltenyi Biotec). The cell suspension was filtered through a 70 µm followed 40 µm cell strainer (SPL Life Sciences), to remove residual debris. After adding PBS containing 0.5% bovine serum albumin (BSA), cells were centrifuged at 200 × g for 8 minutes at 4 °C. The pellet was resuspended in ACK buffer (Thermo Fisher Scientific, #A1049201) to lyse red blood cells, mixed gently, and incubated at room temperature for 3 minutes. Lysis was quenched by adding 5 mL of 0.5% BSA/PBS, followed by centrifugation at 200 × g for 5 minutes at 4 °C. The supernatant was removed, and the pellet was resuspended in 2 mL of 0.5% BSA/PBS and centrifuged again under the same conditions (200 × g, 5 minutes, 4 °C). The final cell pellet was resuspended in 0.5% BSA/PBS and collected as bone marrow cells.

#### Bone lining cell isolation

Bone lining cells were isolated as described^84^. Briefly, after bone marrow cells were flushed out, the bones were roughly chopped into small fragments using scissors and forceps, followed by enzymatic dissociation with Type I collagenase solution (Worthington Biochemical Corporation, #CLS-1 LS004196) supplemented with 20 µL of DNase I (Merck) for 30 minutes at 37°C with gentle agitation. Dissociated cells were filtered through a 70 µm followed 40 µm cell strainer (SPL Life Sciences). After adding PBS containing 0.5% bovine serum albumin (BSA), cells were centrifuged at 300 × g for 5 minutes at 4 °C. The pellet was resuspended in ACK buffer (Thermo Fisher Scientific) and incubated on ice for 5 minutes. Lysis was quenched by adding 25 mL of PBS, and the suspension was passed through a 70 μm cell strainer. Cells were centrifuged again at 1,400 rpm for 5 minutes at 4 °C, and the pellet was resuspended in 0.5% BSA/PBS. The final cell pellet was resuspended in 0.5% BSA/PBS and collected as bone lining cells.

Cell viability was assessed by trypan blue exclusion (Nacalai Tesque, #29853-34), ensuring >90% viability. Bone marrow cells and bone lining cells were pooled separately for each mouse and stained with 7-Aminoactinomycin D (7-AAD, 1:5; BD Biosciences) to sort the living cells, processed immediately for single-cell capture.

#### scRNA-seq library construction and sequencing

Single-cell suspensions were processed using the 10x Genomics Chromium GEM-X Single Cell 3’ Reagent Kit (v4). Approximately 20,000 cells per sample were loaded onto the Chromium Controller, and libraries were generated with unique dual indices (UDIs) according to the manufacturer’s instructions. Libraries were amplified for 13 cycles and quality-checked using the Agilent High Sensitivity DNA kit. cDNA amplification and library construction were performed following GEM-X protocol recommendations. Sequencing was performed on an Illumina NovaSeq X Plus platform, generating 150 bp × 2 paired-end reads with a target depth of 50,000 reads per cell. Raw data were demultiplexed and aligned to the human and mouse reference genome using Cell Ranger (v9.0.0, 10x Genomics) with the reference package *refdata-gex-GRCh38_and_GRCm39-2024-A.tar.gz* (downloaded from the 10x Genomics website).

#### scRNA-seq data processing, integration, and clustering

Genes expressed in fewer than five cells were excluded. Cells with < 200 or > 15,500 detected genes, or mitochondrial gene content >10%, were considered low-quality and removed. After filtering, approximately 11,000-15,000 cells per sample were retained for downstream analysis. Data were normalized and analyzed using Seurat (v5.2.0) in R, including normalization, dimensionality reduction, clustering, and differential gene expression analysis according to the Seurat tutorial (https://satijalab.org/seurat/articles/pbmc3k_tutorial). Datasets from control, Day 10, and Day 20 bone marrow and bone-lining cells were integrated using the *FindIntegrationAnchors* and *IntegrateData* functions with default parameters. The optimal dimensionality was determined using the *ElbowPlot* function, and the first 10 dimensions were selected for integrated data. Graph-based clustering was performed using the *FindNeighbours* and *FindClusters* functions. For differential expression analysis, marker genes for each cluster were identified using the *FindAllMarkers* function (only.pos = TRUE, min.pct = 0.25, logfc.threshold = 0.2, test = “wilcox”). The top 30 genes per cluster were exported for downstream annotation.

Cell type annotation was performed by systematically comparing and validating the top 30 differentially expressed genes of each cluster with reference marker genes curated in the CellMarker 2.0 database (http://www.bio-bigdata.center) and PanglaoDB (https://panglaodb.se). For genes that were not listed in these databases, annotation was supplemented by literature curation through PubMed searches to identify representative marker genes reported in multiple independent studies. In cases where the same cell type was subdivided into multiple clusters, populations were labeled with numerical identifiers (e.g., *Neutrophil-1*, *Neutrophil-2*) to distinguish cluster-specific subgroups. When clusters contained heterogeneous features indicative of more than one cell type, they were annotated as *Mixture* to reflect their mixed cellular composition, typically consisting of marker genes derived from more than one lineage.

### Gene Ontology (GO) enrichment analysis

Gene symbols of differentially expressed genes were converted to Entrez Gene IDs using the *org.Mm.eg.db* package (mapIds, keytype = “SYMBOL”, column = “ENTREZID”, multiVals = “first”). GO enrichment analysis was then performed with *clusterProfiler* (v4.12.0) using the *enrichGO* function against the Biological Process (BP) ontology with the following parameters: OrgDb = org.Mm.eg.db, keyType = “ENTREZID”, pAdjustMethod = “BH”, pvalueCutoff = 0.05, and qvalueCutoff = 0.05. Unless otherwise specified, other arguments were left at package defaults. Significantly enriched GO terms were visualized as dot plots using the *enrichplot* package (dotplot, showCategory = 20). To highlight immune-related categories, enriched terms containing the keyword “Immune” were plotted.

### Trajectory analysis

To reconstruct lineage trajectories, Seurat objects were converted into Monocle3 (v1.4.25) cell data sets (CDS) using the *SeuratWrappers* package. Gene symbols were assigned to the *gene_short_name* column, and cluster annotations obtained from Seurat were transferred into the Monocle3 object. Dimensionality reduction coordinates from Seurat (UMAP) were imported into Monocle3 and used for graph learning.

The trajectory graph was inferred with the *learn_graph* function, and root cells were defined based on prior biological annotation. Cells were ordered along pseudotime using the *order_cells* function. Pseudotime distributions across clusters were visualized by density plots, and the dynamics of selected genes (e.g., *CSF-3* [*G-CSF*]*, Cxcl16*) were evaluated along pseudotime using scatter plots. Branch-associated genes were identified using the *graph_test* function, which detects genes with significant expression changes across trajectory bifurcations. Genes with *q* < 1×10⁻⁴ were considered significant, and the top-ranked candidates were visualized using the *plot_genes_in_pseudotime* function. The list of significantly branched genes was exported for downstream analyses.

### AUCell gene-set analysis

To evaluate the enrichment of tumor-associated macrophage (TAM) subtype–related gene signatures at the single-cell level, AUCell analysis was performed using the AUCell (v1.24.0) package. The normalized RNA expression matrix was extracted from a Seurat object generated from the reclustering of cluster 7 macrophage populations. The Seurat object was normalized using the *NormalizeData* function.

#### Definition of TAM gene signatures

A total of 19 gene sets representing distinct TAM subtypes were curated for AUCell analysis. 6 major TAM-related gene signatures—“Inf_TAM”, “Angio_TAM”, “Lipid-associated_mac (LAM)”, “Reg_TAM”, “IFN_TAM” and “Prolif_TAM”—were custom-defined based on comprehensive integration of multiple published datasets^21,85,86,39,36, 37, 40–44, 87–90^. These custom signatures were established by integrating representative marker genes that consistently characterized the indicated TAM phenotypes across studies, while excluding genes with low or undetectable expression in our dataset to minimize technical noise and improve the robustness of the analysis.

In addition, several TAM signatures were adopted directly from previously published reference datasets, including “CXCL8+_monocyte”, “M1_macrophage”, “FABP4+_macrophage”, “CD14+_monocyte”, “M2_TAM” ^91^ ; and “Angiogenesis”, “Complement”, “Immune_suppression”, “Antigen_processing_and_presentation” ^92^ ; and “Mac_S4_pro-inflammatory” and “Mac_S2_anti-inflammatory”^93^; and “Lipid_Mo_TAM” and “Transitory_Mo_TAM”^34^. Genes with low or undetectable expression in our dataset were excluded to minimize technical noise and improve the robustness of the analysis.

#### Definition of non-tumor disease-associated macrophage gene signatures

A total of 17 gene sets representing distinct non-tumor disease-associated macrophage-related gene signature was curated for AUCell analysis. 3 major gene signatures— “Bacterial_infection”, “Virus_infection”, and “Autoimmunity”—were custom-defined based on comprehensive integration of multiple published datasets^39, 48–58^. These custom gene sets were designed by integrating representative marker genes that consistently defined the indicated macrophage populations across species and experimental contexts. All non-tumor disease-associated macrophage-related gene sets were cross-validated for consistency among studies, and genes with low or undetectable expression in our dataset were excluded before AUCell analysis.

Several gene sets were directly adopted from published datasets without modification, including senescence-associate gene sets (“G2M_phase”, “Surveying”, “Stressed”, “Poised_Inflammatory”, “Baseline_surfactant_turnover”) ^45^, osteoarthritis-associated gene sets (“Crip1hiCav1+”, “Trem2hi_Fcrls+”, “Lyve1hiFolr2hi”, “IFN-R_Mono-Mac”, “MHCllhi”) ^46^, and bone fracture-associated gene sets (“Cluster_29”, “Cluster_16”, “Cluster_10”, “Cluster_12”)^47^. Genes with low or undetectable expression in our dataset were excluded before AUCell analysis.

#### Definition of TAM gene signatures from breast cancer metastases in lung and brain and from primary tumors

Gene sets were directly adopted from published datasets without modification, including breast cancer lung metastasis–associated TAM gene sets (“interstitial macrophage [IM]_Proliferative,” “IM_Crip1,” “alveolar macrophage [AM]_Inflammatory,” “AM_Lipid,” “AM_Homeostatic,” “AM_Antigen,” “IM_Antigen,” and “IM_Mrc1”)^65^ ; additional lung metastasis–associated TAM gene sets (“Mac_Isg20” and “Mac_Cd81”)^66^, ; non–small-cell lung cancer (NSCLC)–associated TAM gene sets (“Infl_monocyte,” and “Monocyte_mac [MoMac]”)^68^, ; breast cancer brain metastasis–associated TAM gene sets (“Microglia” and “pvMacrophage”)^67^, ; and glioma-associated TAM gene sets (“Macrophage” and “MG7”)^69^. Genes with low or undetectable expression in our dataset were excluded before AUCell analysis.

#### Establishment of a breast cancer PDX model with spontaneous lung metastasis and identification of TAM gene signatures by scRNA-seq

Primary breast cancer tissues were minced into ∼1-mm pieces and orthotopically implanted into the mammary fat pads of immunodeficient mice (4–5 tissue pieces per site, two sites per mouse, one on each side) as described^16^. Primary mammary tumors developed approximately 4–5 months after transplantation and were surgically excised once they reached a palpable size.

Spontaneous lung metastases appeared 2–3 months after removal of the primary tumors. The lungs containing metastatic nodules were resected from tumor-bearing mice. The samples were briefly washed with PBS and minced into small fragments. The tissue fragments were enzymatically dissociated with collagenase (Merck), dispase (1 U/mL; STEMCELL Technologies, #07913) and DNase I (100 U/mL; Merck) at 37 °C with gentle agitation. Cell suspensions were triturated, passed through 70-μm strainers, and treated with red blood cell lysis buffer (Merck). Single-cell RNA sequencing (scRNA-seq) was performed using dissociated single-cell suspensions from metastatic lung tissues.

Single-cell suspensions were processed using the 10x Genomics Chromium NEXT GEM Single Cell 3’ Reagent Kits v3.1 (Dual Index). Approximately 10,000 cells per sample were loaded onto the Chromium Controller, and libraries were generated with unique dual indices (UDIs) according to the manufacturer’s instructions. Libraries were amplified for 13 cycles and quality-checked using the Agilent High Sensitivity DNA kit. cDNA amplification and library construction were performed following protocol recommendations. Sequencing was performed on an Illumina NovaSeq X Plus platform, generating 150 bp × 2 paired-end reads with a target depth of 50,000 reads per cell. Raw data were demultiplexed and aligned to the human and mouse reference genome using Cell Ranger (v9.0.0, 10x Genomics) with the reference package *refdata-gex-GRCh38_and_GRCm39-2024-A.tar.gz* (downloaded from the 10x Genomics website).

The scRNA-seq data were processed and clustered into 15 distinct clusters (**Extended Data Fig. 8a,b**). Among these, two macrophage clusters were identified-patient 10 TAM1 (P10_TAM1) and patient 10 TAM2 (P10_TAM2). The top 20 differentially expressed genes (DEGs) for each macrophage cluster were extracted (**Supplementary Tables 10 and 11**) and defined as two macrophage-specific gene signatures, designated as lung metastasis “P10_TAM1” and “P10_TAM2,” respectively. These gene signatures were subsequently used for gene signature analysis.

#### Definition of NF-κB gene signatures

To evaluate the activation status of the NF-κB signaling pathway within macrophage subsets, we generated a custom NF-κB activation gene set based on canonical pathway components and downstream target genes^94, 95^. This gene set included key transcriptional regulators and effector genes involved in the NF-κB signaling cascade: *Rela*, *Relb*, *Rel*, *Nfkb1*, *Nfkb2*, *Nfkbia*, *Nfkbiz*, *Chuk*, *Ikbkb*, *Ikbke*, *Tnfaip3*, *Ccl2*, *Il6*, *Tnf*, *Cxcl1*, *Ptgs2*, *Nos2*, and *Birc3*. The selection was based on genes broadly recognized as mediators or readouts of classical NF-κB activation in macrophages. AUCell scoring of this gene set enabled comparative assessment of NF-κB pathway activity across macrophage clusters.

#### Heatmap and rose diagram visualization of AUCell scoring

Gene ranking matrices were constructed using the *AUCell_buildRankings* function, and the area under the curve (AUC) for each gene set was calculated using *AUCell_calcAUC*. The resulting AUC matrix was transposed and incorporated into the Seurat metadata. Cluster-level average AUCell scores were computed by grouping cells according to Seurat cluster identity and calculating mean AUC scores for each gene set. The averaged matrix was row-wise z-score–normalized to obtain relative enrichment values. Heatmap visualization was performed using the ComplexHeatmap (v2.20.0) package. A diverging color scale (blue– white–red) was generated using *circlize::colorRamp2* across a value range of –2.5 to +2.5, representing relatively low to high AUCell scores. Rows (gene sets) were hierarchically clustered, whereas columns (clusters) were ordered as pseudotime.

To further illustrate relative enrichment patterns across clusters, Rose plots were generated based on the cluster-averaged relative AUCell scores obtained from the heatmap. For each gene set, the relative AUCell scores were scaled by Z-score normalization, and the resulting values were used to determine the sector length in the Rose plot. Visualization was performed using the ggplot2 (v3.5.2) package with a polar coordinate system, and color gradients consistent with the heatmap were applied to indicate the relative AUCell (Z) values.

#### Correlation analysis of macrophage-related gene signatures

To examine the transcriptional relationships among distinct macrophage programs, AUCell scores were calculated from normalized RNA expression values of macrophage subclusters using the *AUCell* package. Cell-wise gene rankings were generated with *AUCell_buildRankings*, and AUCell scores were computed for each curated gene-set collection using *AUCell_calcAUC*.

Three categories of curated macrophage-related gene sets were analyzed: 19 TAM gene sets, 17 non-tumor disease–associated macrophage gene sets, and 20 metastasis-associated TAM gene sets derived from breast cancer metastases in the lung and brain, as well as from primary non–small-cell lung cancer and glioma.

Pairwise Pearson correlation coefficients were computed between AUCell scores from these three categories to evaluate similarities in transcriptional activity patterns across macrophage subtypes. Three pairwise comparisons were performed: (a) TAM vs. non-tumor disease–associated macrophage gene sets, (b) TAM vs. metastasis-associated TAM gene sets, and (c) non-tumor disease–associated vs. metastasis-associated TAM gene sets. The resulting matrices were visualized as clustered heatmaps with the *ComplexHeatmap* package, where color intensity represents the Pearson correlation coefficient (r) ranging from –1 to 1. Hierarchical clustering was performed on both rows and columns to group gene sets with similar correlation profiles.

### Statistics

Statistical analyses were performed in R environment (v4.4.3) or GraphPad Prism software (v10.5.0). Statistical methods and details are described in the figure legends. Briefly, for normally distributed data, the significance was calculated with the unpaired, 2-tailed Student’s *t* test. Comparisons between more than 2 groups was performed using 1-way ANOVA or 2-way ANOVA with Dunnett’s post hoc test or Bonferroni’s post hoc test. The in vivo tumorigenesis data were analyzed by 2-way ANOVA, followed by post hoc multiple comparisons adjusted using the two-stage linear step-up procedure of Benjamini, Krieger, and Yekutieli. The correlation between variables was evaluated using the Pearson correlation test in GraphPad Prism. The Pearson correlation coefficient (r) and corresponding *p*-values were computed to assess the strength and significance of linear associations. Statistical analyses were performed in R using RStudio. AUCell scores among macrophage clusters were compared using one-way ANOVA followed by Bonferroni’s post hoc test. p < 0.05 was considered statistically significant.

## Data availability

The RNA sequencing (RNA-seq) and single-cell RNA sequencing (scRNA-seq) data generated in this study are available from the DNA Data Bank of Japan (DDBJ) Sequence Read Archive (DRA). Bulk RNA-seq datasets are deposited under accession numbers PRJDB39673 and PRJDB39682. Single-cell RNA-seq datasets are deposited under accession numbers PRJDB39704 and PRJDB39737. Processed gene-expression matrices and cell-level metadata used for downstream analyses are available from the corresponding author upon reasonable request. Source data underlying all main and extended data figures are provided with this paper.

## Acknowledgements

We thank Professor J. Massagué for his critical feedback on this manuscript. This work was supported by JST SPRING, Grant Number JPMJSM2135 to H.Z. This work was supported in part by JSPS KAKENHI Grant Numbers JP22H04923 (CoBiA), JP21H02761, JP 22H02761, JP 23K18237, JP 24K02304, JP 25K22575 to N.G. and AMED Project for Cancer Research and Therapeutic Evolution JP18ck0106194 to N.G. and for Promotion of Cancer Research and Therapeutic Evolution [P-PROMOTE] (JP22ama221517 and JP25ama221610 to E.A.S., JP24ama22114, JP24ama221539, JP25ama22114, and JP25ama221539 to N.G.). This work was supported in part by Grants-in-Aid from Nakatani Foundation to E.A.S. and H.S. This work was supported in part by Grants-in-Aid from the Uehara Memorial Foundation, Princess Takamatsu Cancer Research Fund (PTCRF), Takeda Science Foundation and The Yasuda Medical Foundation to N.G.

## Author contributions

YT and HZ performed most of the experiments, analyzed the data, carried out the bioinformatics analyses, and contributed to manuscript writing. TM performed additional experiments and data analyses. TH provided the luciferase reporter–expressing lentiviral vector and contributed to discussions. KI, KH, and SI performed RNA sequencing and data analysis. MY, MT, YK, HT, NI, and K-IT provided clinical samples. EAS provided the protocol for immunofluorescence of patient-derived organoids. EH provided mCherry-expressing CAFs. S-I T and MH performed bioinformatics analyses. HW and KO contributed to expert discussions. SS and YM provided pathological specimens. FA and TS provided the protocol for isolating bone-lining cells and contributed to expert discussions. AT supervised the study. KO performed single-cell analyses and contributed to expert discussions. HC provided the protocol for culturing patient-derived breast-cancer organoids and contributed to expert discussions. TT contributed important conceptual insight and bioinformatics expertise. NG conceptualized the study, designed the research, and drafted and finalized the manuscript.

