## Extended.data.figures1-9and legends for "Highly plastic macrophage niches orchestrate acquired quiescence and reactivation in breast-cancer bone metastasis"

### Extended Data Fig. 1

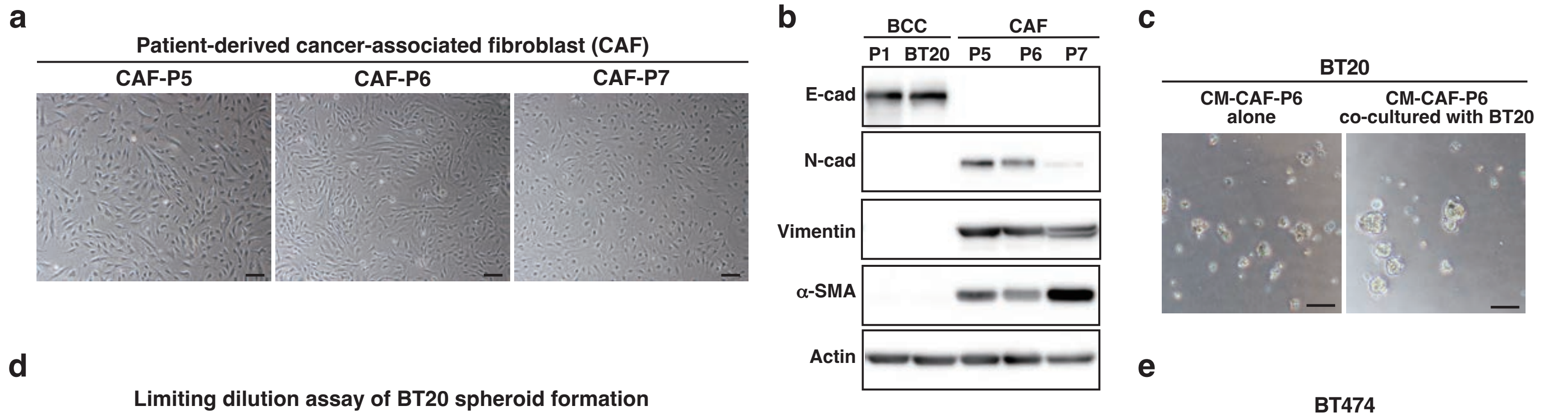


**d**

Limiting dilution assay of BT20 spheroid formation

| Cell number | 125 | 250 | 500 | 1000 | 2000 | 4000 | Tumor-initiating cell frequency | P-value |
|---|---|---|---|---|---|---|---|---|
| CM-CAF-P6 alone | 0/8 | 0/8 | 0/8 | 2/8 | 5/8 | 8/8 | 1/699 | 0.004 |
| CM-CAF-P6 co-cultured with BT20 | 0/8 | 1/8 | 2/8 | 4/8 | 8/8 | 8/8 | 1/258 | |

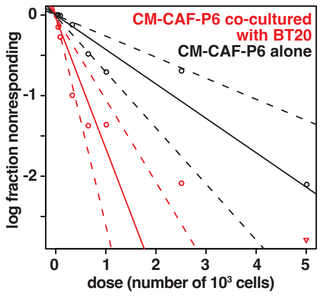


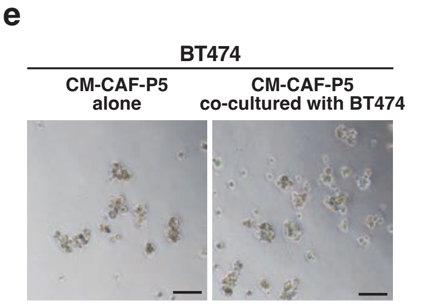


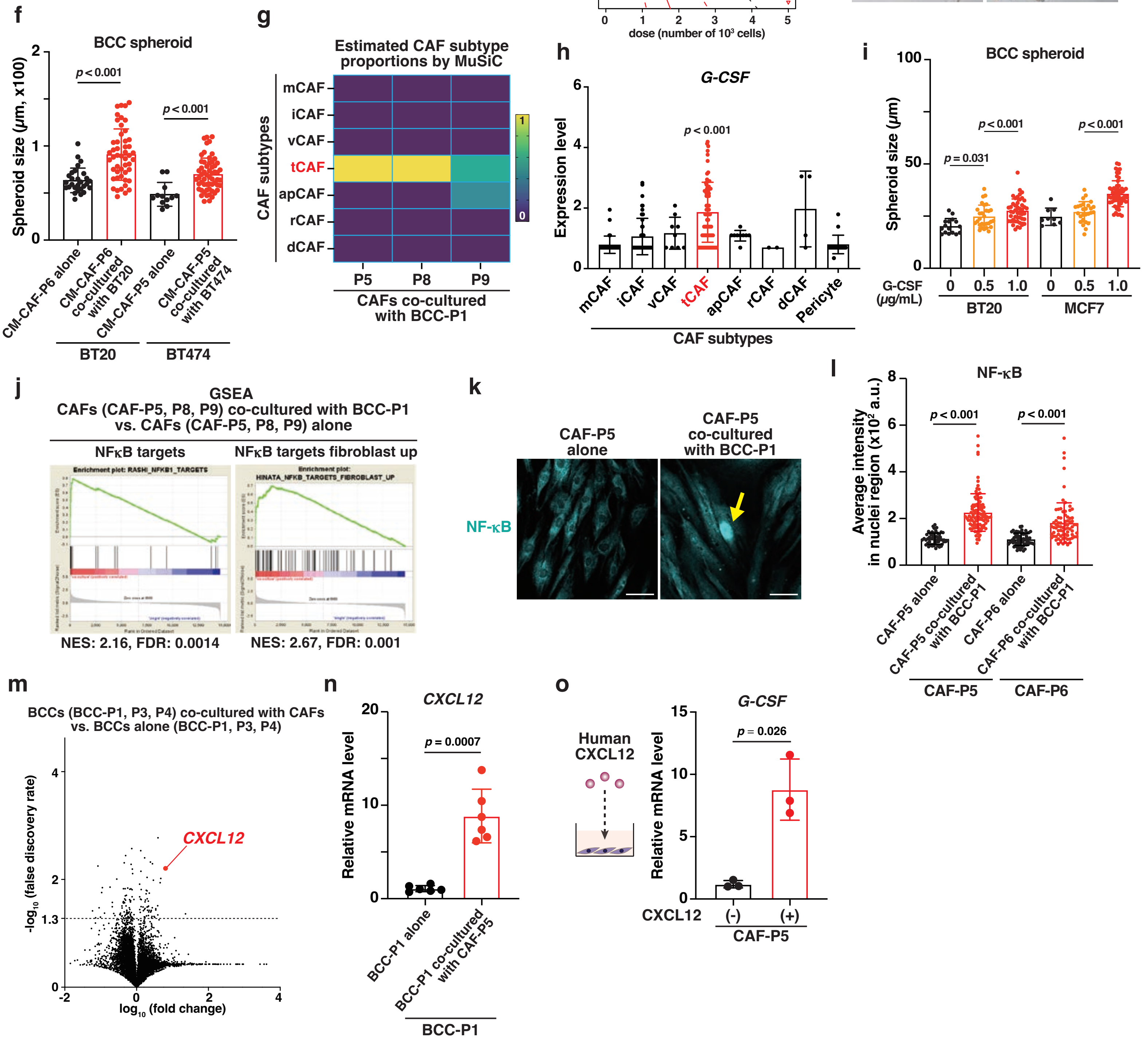

### Extended Data Fig. 2

**a**

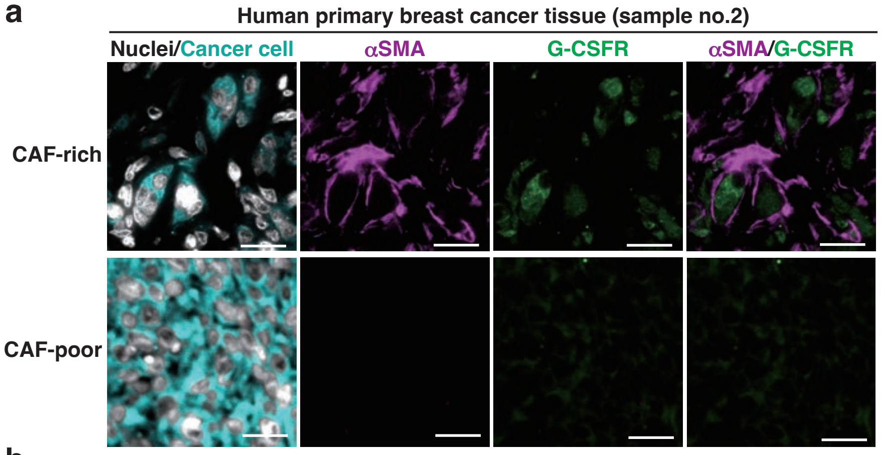


**b**

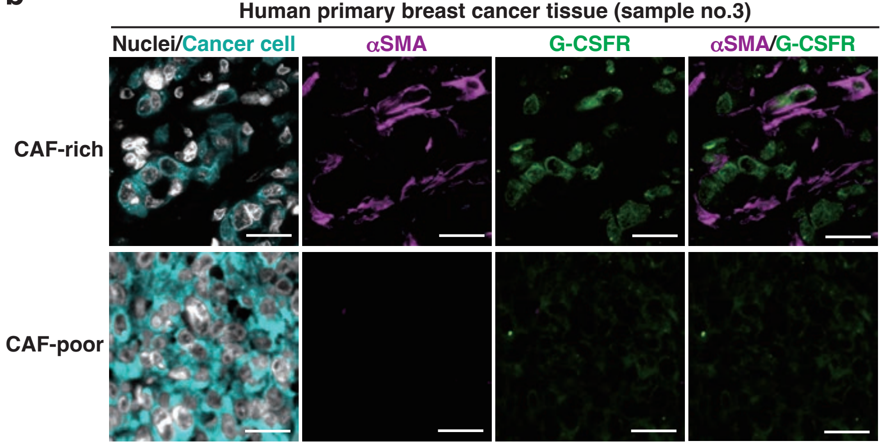


**c**

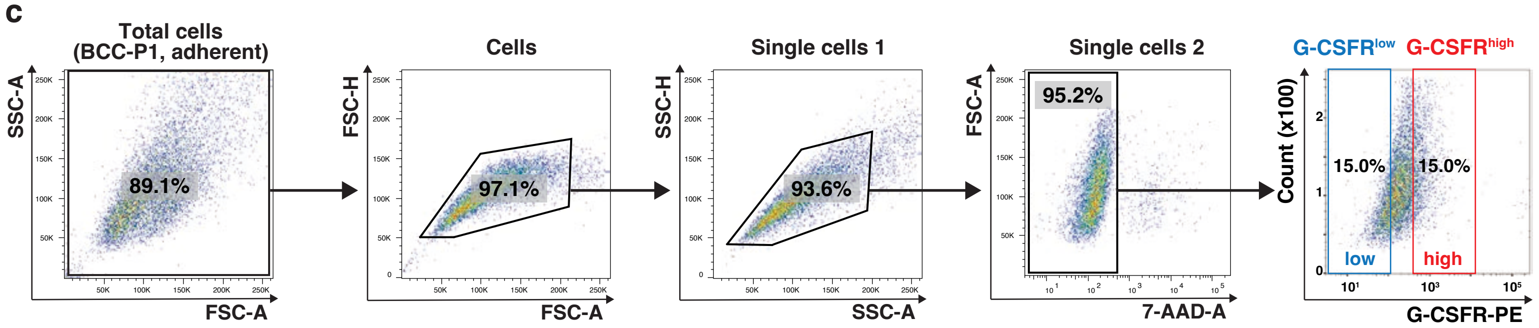


**d**

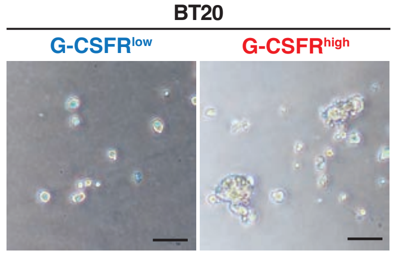


**e**

Limiting dilution assay of BT20 spheroid formation

| Cell number | 6 | 13 | 25 | 50 | 100 | 200 | Tumor-initiating cell frequency | P-value |
|---|---|---|---|---|---|---|---|---|
| G-CSFR$^{low}$ | 0/6 | 0/6 | 0/6 | 1/6 | 2/6 | 3/6 | 1/344.5 | 0.0252 |
| G-CSFR$^{high}$ | 0/6 | 0/6 | 1/6 | 3/6 | 5/6 | 6/6 | 1/85.7 | |

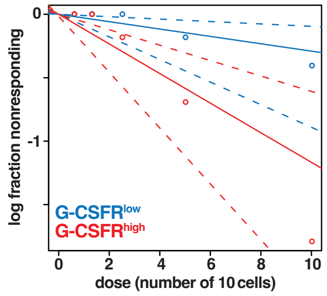


**f**

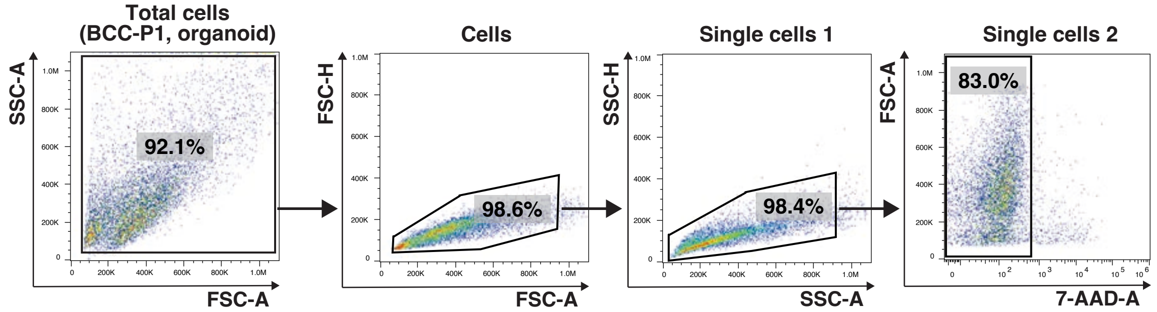


**g**

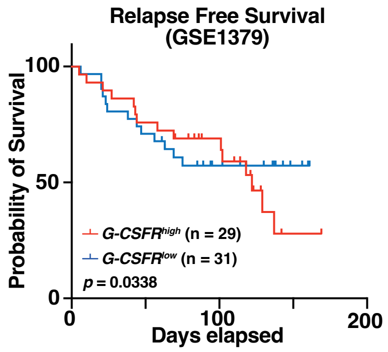

#### Extended Data Fig. 3

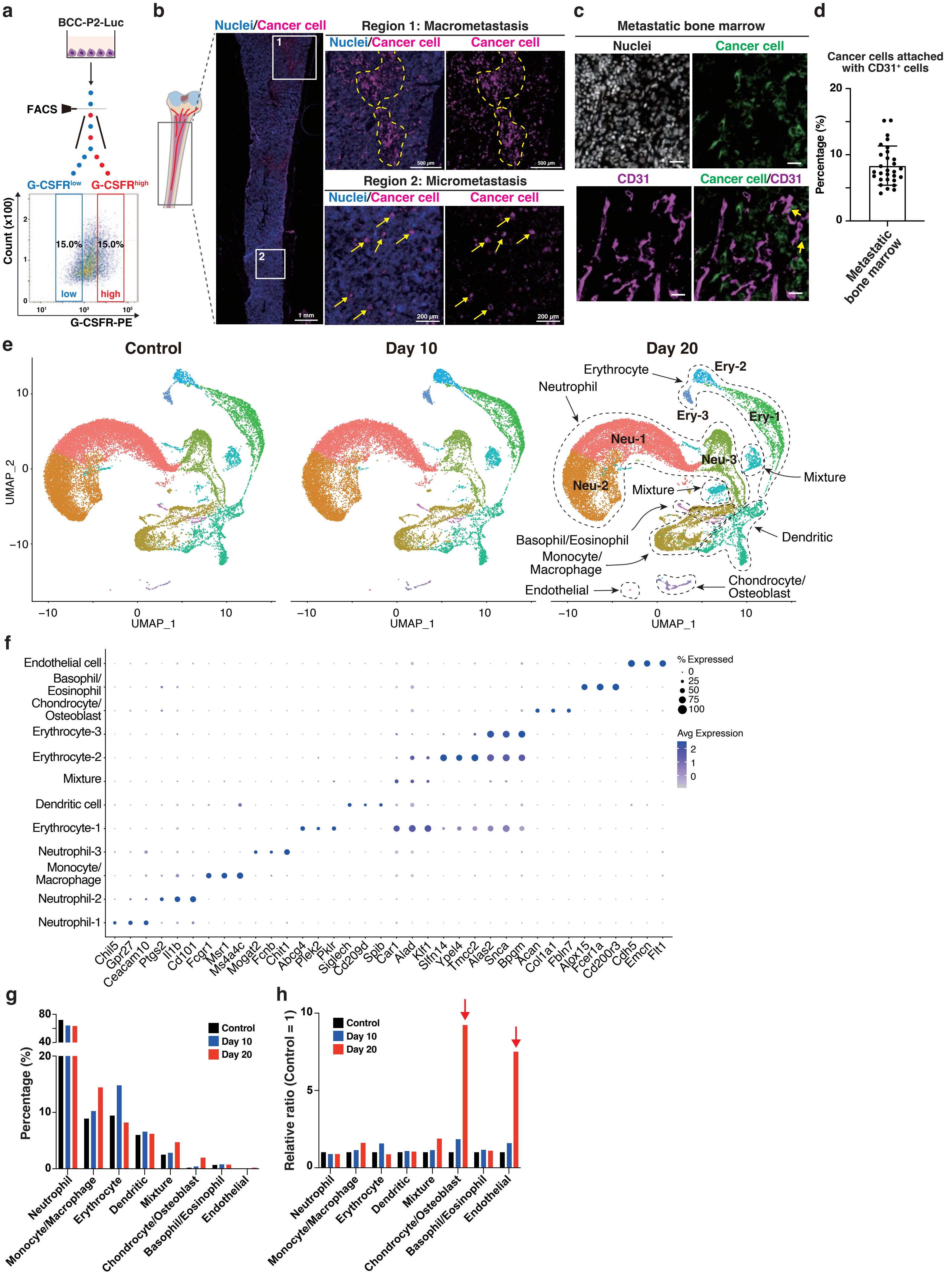

#### Extended Data Fig. 4

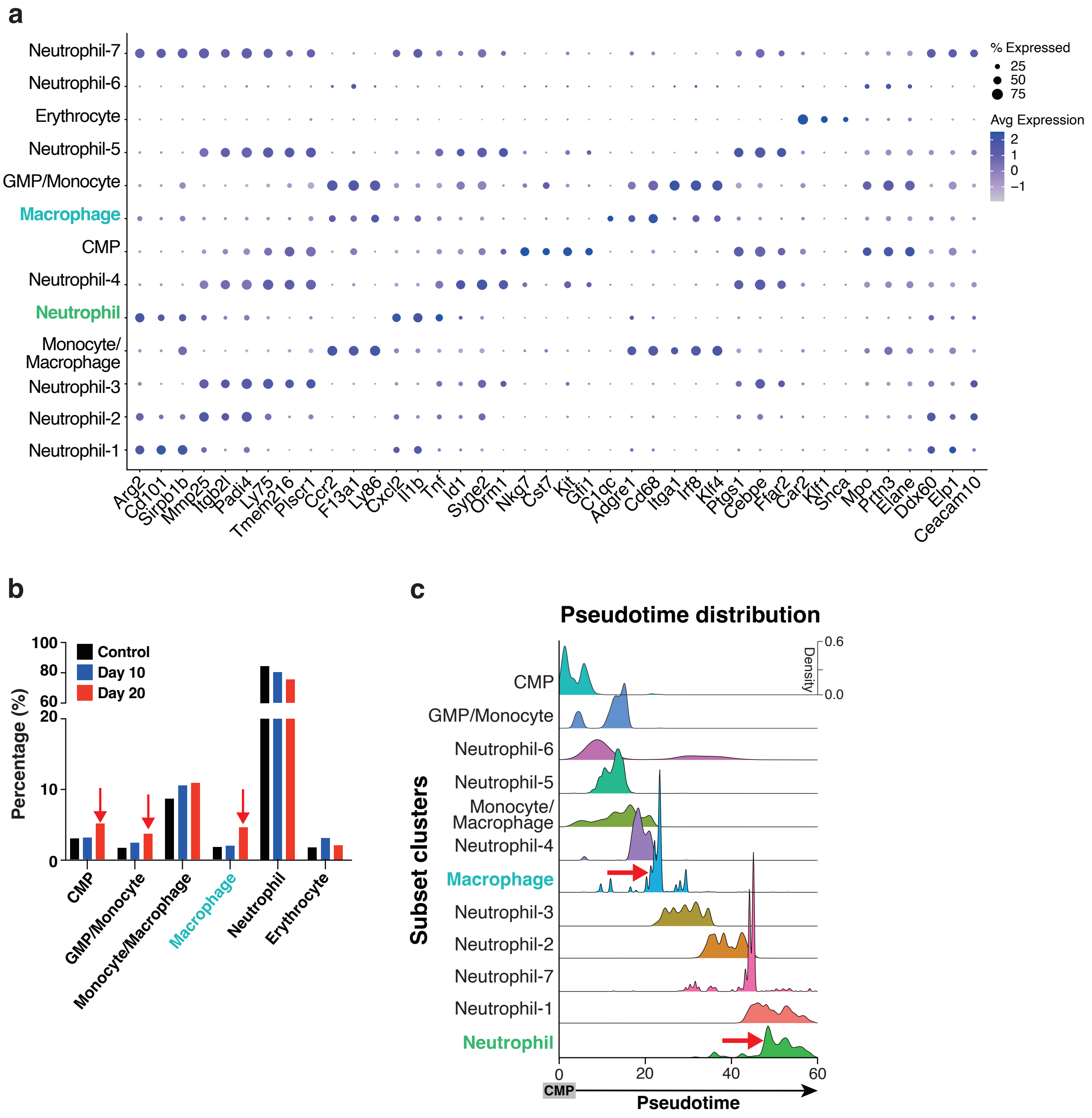

#### Extended Data Fig. 5

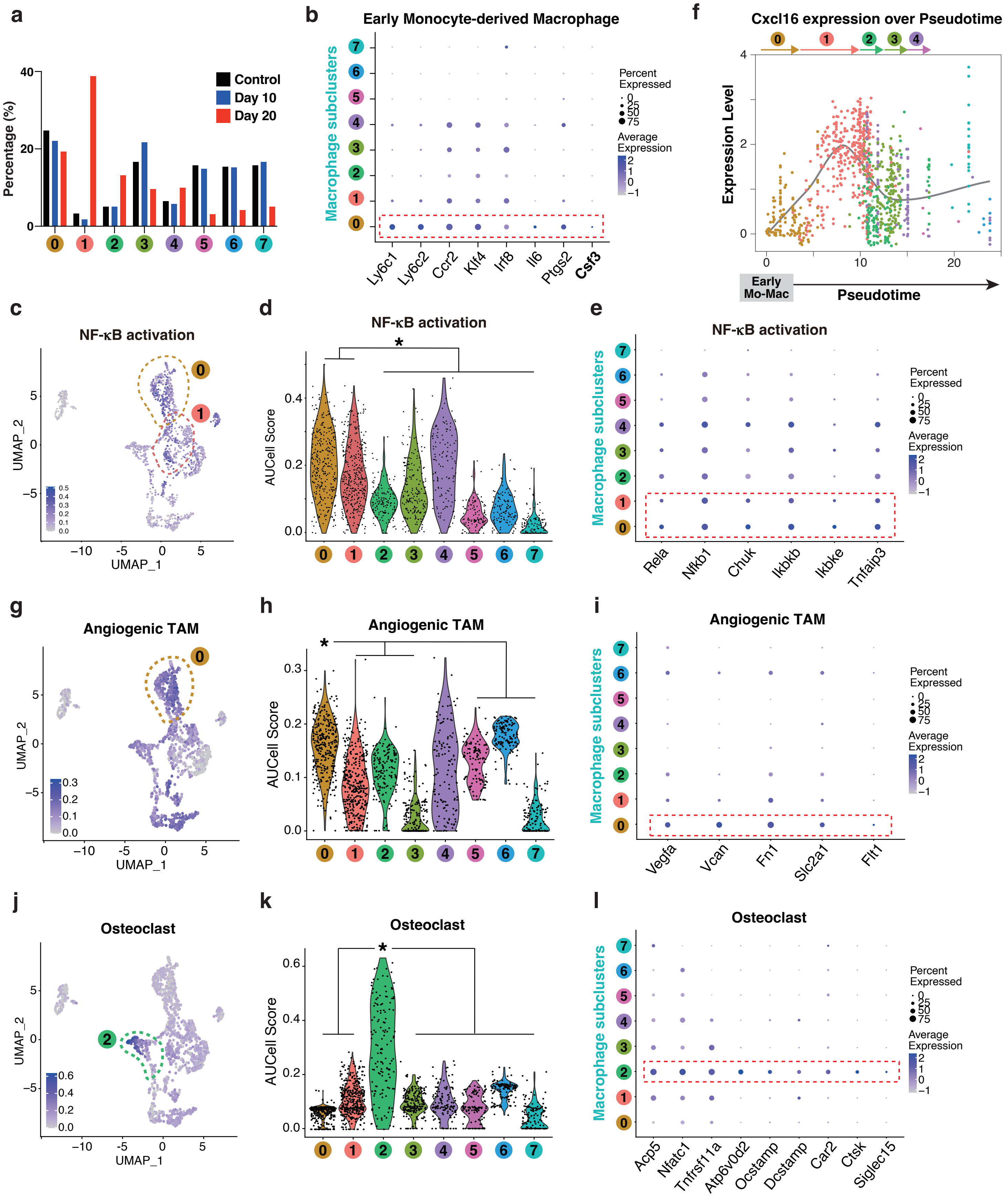

#### Extended Data Fig. 6

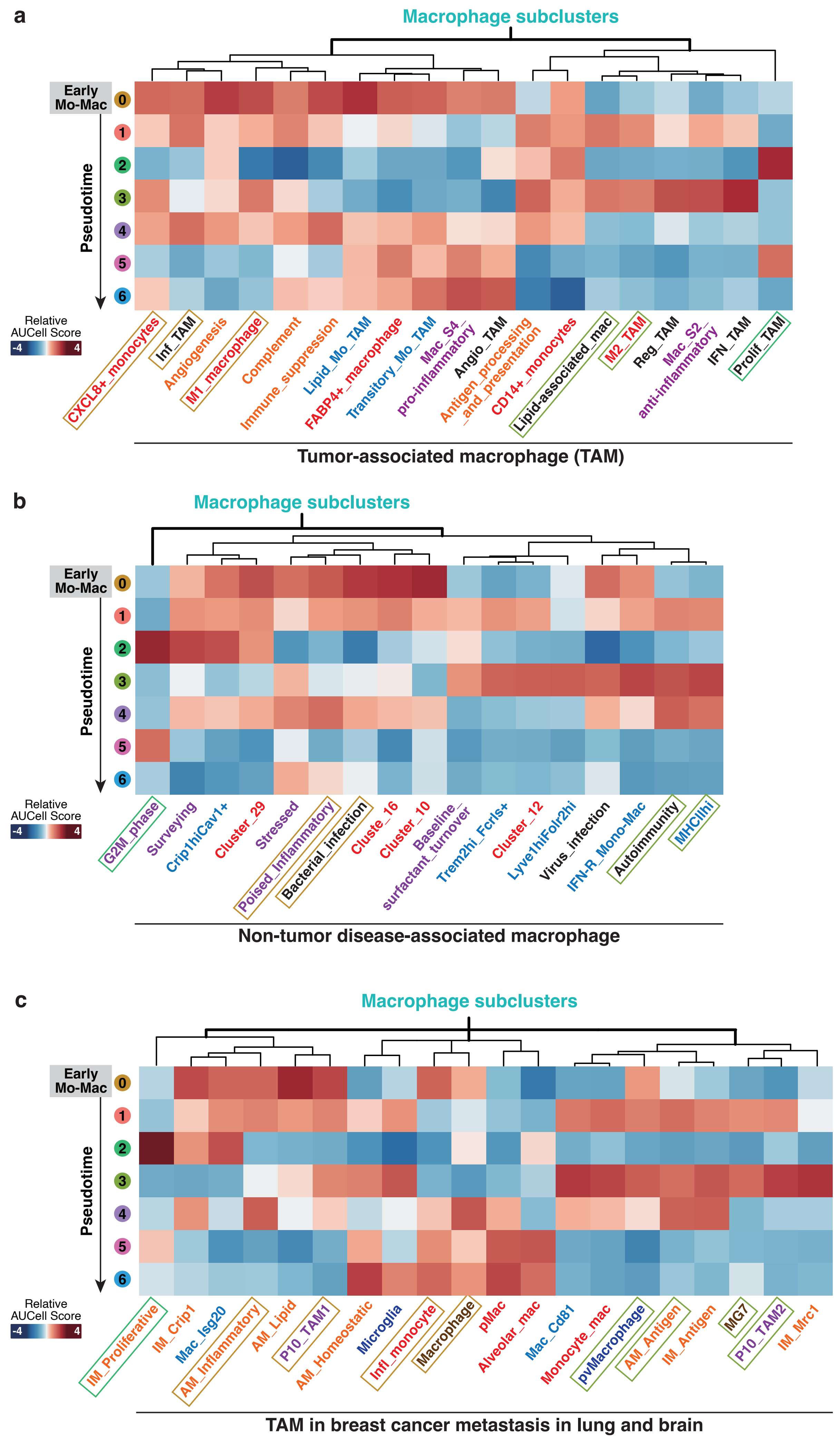

#### Extended Data Fig. 7

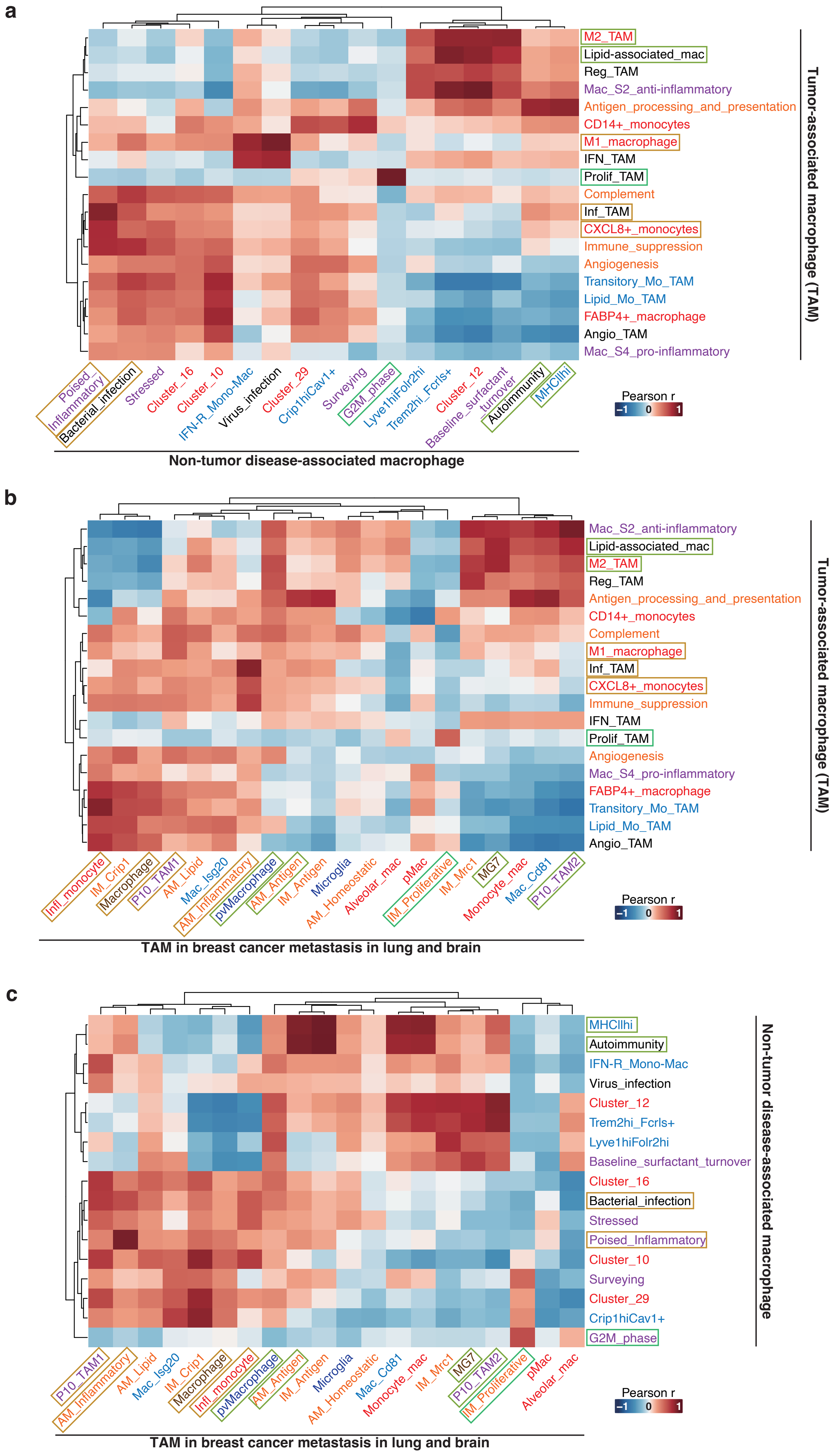

#### Extended Data Fig. 8

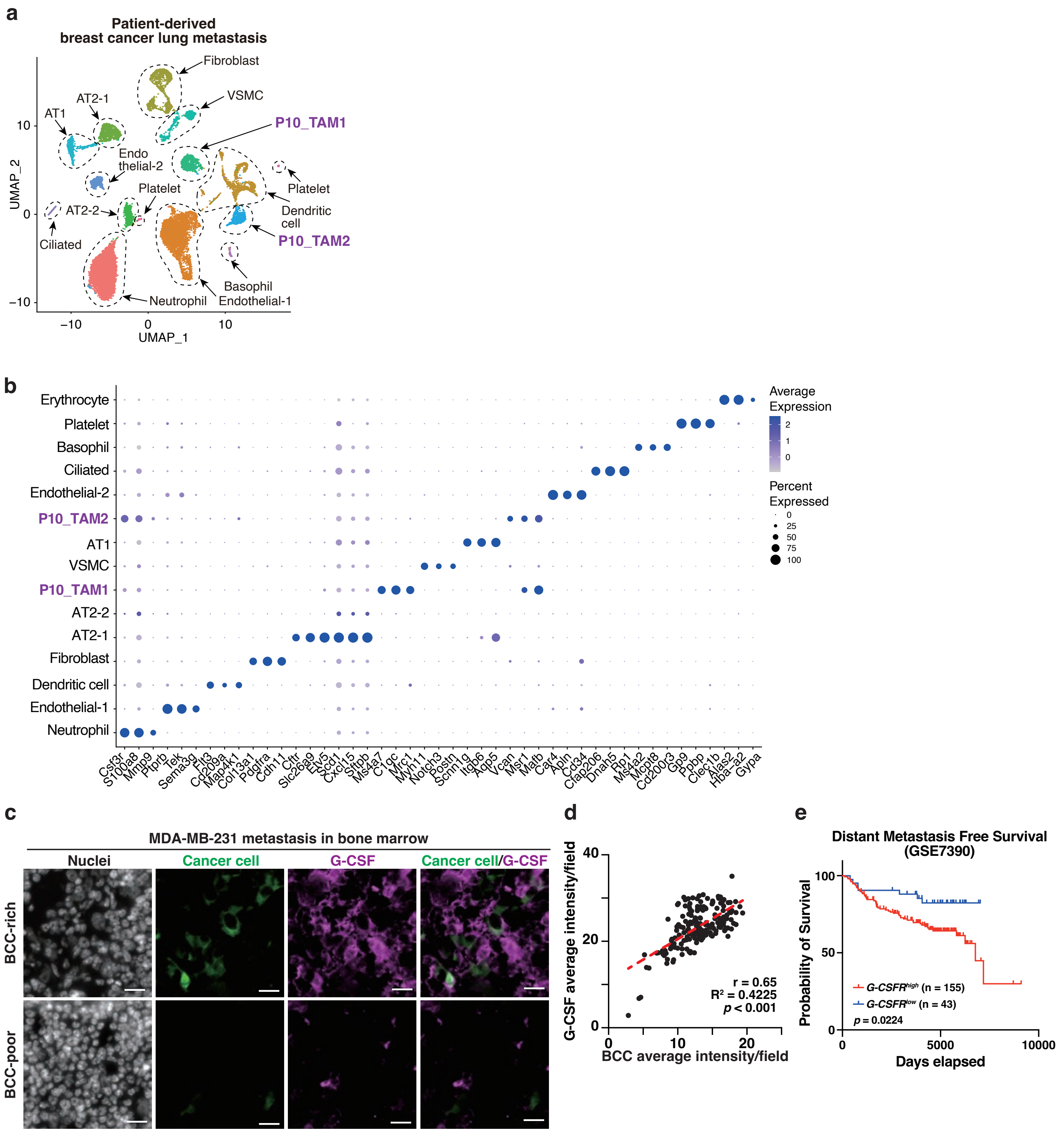

### Extended Data Fig. 9

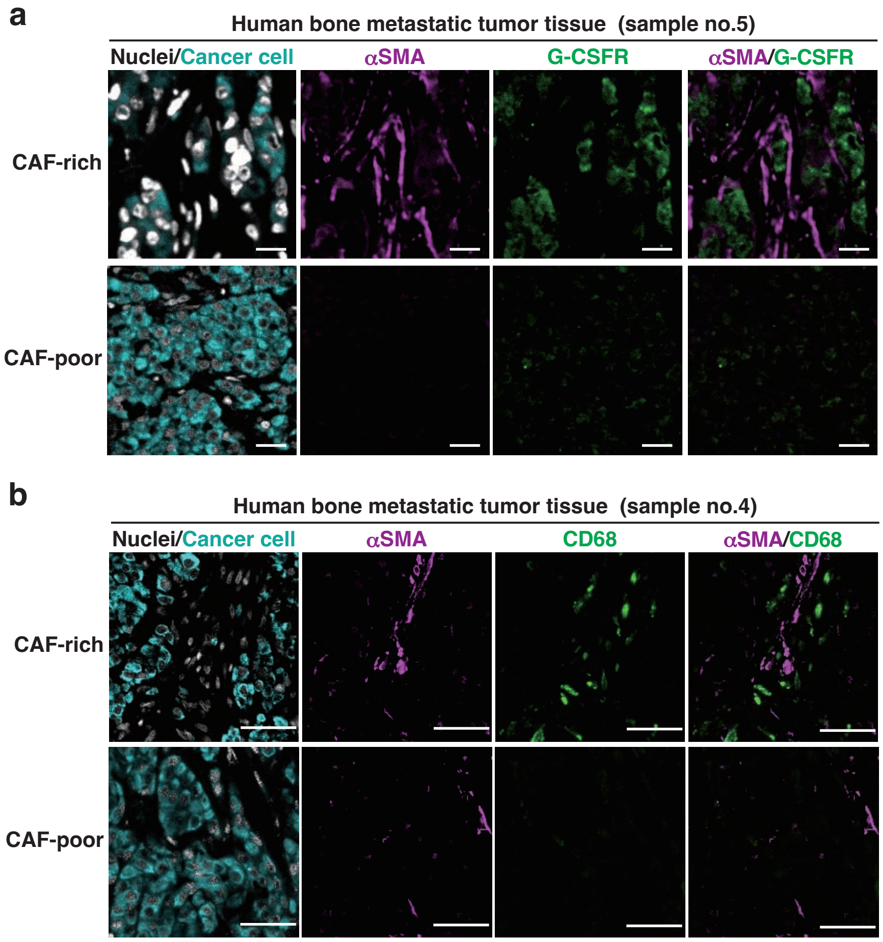

#### Extended Data Fig legends

##### Extended Data Fig. 1

**Patient-derived tumor-like CAFs (tCAFs) produce G-CSF via NF-κB signaling triggered by CXCL12 from breast cancer cells (BCCs) through paracrine interactions.**

**a,** Representative bright-field images of patient-derived CAFs (P5, P6, P7). Scale bars, 100 μm. **b,** Whole-cell lysates from BCCs (BCC-P1, BT-20) and CAFs (P5, P6, P7) immunoblotted with the indicated antibodies. E-cad, E-cadherin; N-cad, N-cadherin. β-Actin served as a loading control. Representative blots were confirmed by at least three independent experiments. **c**–**f,** Representative bright-field images (**c**, **e**), results of the extreme limiting dilution assay (ELDA) (**d**) and quantification of spheroid size (**f**) incubated with conditioned medium (CM) from CAF cultures alone or CAF–BCC co-cultures. Scale bars, 100 μm. **g,** MuSiC deconvolution identified the patient-derived CAFs as tumor-like CAFs (tCAFs). mCAF, myofibroblastic CAF; iCAF, inflammatory CAF; vCAF, vascular CAF; apCAF, antigen-presenting CAF; rCAF, remodeling CAF; dCAF, developmental-like CAF. **h,** Public scRNA-seq datasets revealed that *G-CSF* expression was highest in tCAFs compared with other CAF subtypes (Kruskal–Wallis with Dunn's test). **i,** Quantification of BT20 or MCF7 spheroid size cultured with or without recombinant G-CSF (0.5 or 1.0 μg/ml for 7 days). Data are mean ± SD. Statistical significance was determined by one-way ANOVA with post hoc multiple-comparisons test. **j,** Gene set enrichment analysis (GSEA) comparing CAFs co-cultured with BCCs versus monoculture; gene sets enriched under co-culture are shown (NES, FDR q-values indicated). NES, normalized enrichment score; FDR, false discovery rate. **k,** Representative immunofluorescence images showing nuclear NF-κB p65 in CAFs cultured alone or co-cultured with BCCs. Arrows indicate nuclear p65-positive cells. Scale bars, 100 μm. **l,** Quantification of nuclear NF-κB p65 fluorescence intensities. a.u., arbitrary units. **m–o,** Volcano plot comparing BCCs co-cultured with CAFs versus monoculture (**m**), qPCR analysis of *CXCL12* mRNA expression in BCCs (**n**), and *G-CSF* mRNA expression in CAFs stimulated with recombinant CXCL12 (**o**). Values were normalized to *β-Actin* (*ACTB*). **f**, **l, n**, and **o**, Statistical significance was determined by unpaired, 2-tailed Student's test. Data are mean ± SD.

**Extended Data Fig. 2**

**G-CSF–producing CAFs provide a niche that induces G-CSFR expression and spheroid-forming activity in BCCs.**

**a**, **b,** Representative immunofluorescence images of human breast cancer tissues (sample Nos. 2 and 3) stained for cancer cells (STEM121, human-specific), α-SMA and G-CSFR. Scale bars, 100 µm. **c,** Flow cytometric sorting strategy for breast cancer cells (BCC-P1). **d, e,** Representative bright-field images of tumor spheroids (**d**) and results of the extreme limiting dilution assay (ELDA) (**e**) between G-CSFR$^{low}$ and G-CSFR$^{high}$ BT-20 cell populations. **f,** Flow cytometric sorting strategy for BCC-P1 organoids. **g,** Kaplan–Meier analysis in breast cancer patients (GSE1379) stratified by *G-CSFR* mRNA expression.

**Extended Data Fig. 3**

**Dynamic changes in bone marrow cells are detected by scRNA-seq analysis.**

**a,** Schematic representation of FACS-based isolation of G-CSFR$^{high}$ and G-CSFR$^{low}$ populations from BCC-P2-Luc cells. **b,** Representative immunofluorescence images of femurs at 20 days after caudal artery injection of MDA-MB-231 cells, stained for human-specific cancer cells (STEM121). Nuclei were counterstained with Hoechst. Boxed areas in the left panel are magnified on the right (Region 1, macrometastasis; Region 2, micrometastasis). Dotted circles indicate focal clusters of cancer cells (top), and arrows indicate single disseminated cells (bottom). Scale bars, 1 mm (left), 500 µm (top), and 200 µm (bottom). **c,** Representative immunofluorescence images of femurs at 20 days after caudal artery injection stained for CD31 (endothelial cells) and STEM121 (human cancer cells). Nuclei were counterstained with Hoechst. Scale bars, 100 µm. **d,** Quantification of cancer cells attached to $CD31^+$ endothelial cells. Data are presented as mean ± SD (n = 29 random fields). **e,** Uniform manifold approximation and projection (UMAP) visualization of scRNA-seq data from total bone marrow and bone-lining cells collected from control mice and from 10 and 20 days after caudal artery injection. Cells are colored by cluster identity (n = 2 mice for control, and n = 3 mice each for Day 10 and Day 20). **f,** Dot plot of canonical marker-gene expression across the clusters shown in **e**. Dot color represents scaled average expression, and dot size indicates the proportion of cells expressing each gene. **g**, **h,** Relative abundance of major cell populations across conditions. Fraction of each population among all cells (**g**) and its relative ratio versus the corresponding control (**h**) are shown.

**Extended Data Fig. 4**

**Reclustering of myeloid cells uncovers a previously unrecognized macrophage cluster.**

**a,** Dot plot showing canonical marker-gene expression across subset clusters corresponding to the UMAP in **Fig. 4e**. Clustering of scRNA-seq data based on using marker-gene profiles delineated distinct cell types. Dot color represents scaled average expression, and dot size indicates the fraction of cells expressing each gene within a cluster. **b,** Relative abundance of each subset population across experimental conditions shown in **Fig. 4f. c,** Pseudotime distribution of subset clusters illustrating their temporal progression during metastatic initiation.

**Extended Data Fig. 5**

**NF-κB activation and TAM gene signatures delineate macrophage heterogeneity during bone metastasis initiation.**

**a,** Relative abundance of each macrophage subset across shown in **Fig. 5b. b,** Dot plot showing canonical markers of early monocyte-derived macrophages identified in the UMAP in **Fig. 5a. c**, Feature plot displaying AUCell scores of the NF-κB activation gene signature across macrophage subclusters in the UMAP. **d**, Violin plots showing AUCell scores of the NF-κB activation gene signature across macrophage subclusters. **e,** Dot plot showing expression of NF-kB activation signature genes across the UMAP-defined clusters in **Fig. 5a. f,** *Cxcl16* expression along pseudotime across macrophage subclusters. **g,** Feature plot showing AUCell scores of the angiogenic TAM gene signature across macrophage subclusters in the UMAP. **h,** Violin plots showing AUCell scores of the angiogenic TAM signature across macrophage subclusters**. i,** Dot plot showing expression of angiogenic TAM -relaged genes across the UMAP-defined clusters in **Fig. 5a. j,** Feature plot showing AUCell scores of the osteoclast gene signature across macrophage subclusters in the UMAP. **k,** Violin plots showing AUCell scores of the osteoclast signature across macrophage subclusters**. l,** Dot plot showing expression of osteoclast-related genes across UMAP-defined clusters in **Fig. 5a. d, h, and k,** Statistical significance was determined by one-way ANOVA with Bonferroni's post hoc test.

**Extended Data Fig. 6**

**Heatmap of macrophage subset–signature enrichment reveals high plasticity in newly**

**emerging macrophages along pseudotime.**

**a,** Heatmap showing relative AUCell-based enrichment scores for 19 TAM gene signatures (columns) across macrophage subclusters (rows). **b,** Heatmap showing relative AUCell-based enrichment scores for 17 non-tumor disease-associated macrophage gene signatures (columns) across macrophage subclusters (rows). **c,** Heatmap showing relative AUCell-based enrichment scores of 20 TAM gene signatures derived from breast-cancer metastases in lung and brain (columns) across macrophage subclusters (rows). **a, b, and c**, The color of each TAM gene set corresponds to the reference study from which the signature was derived. Color gradients from blue to red indicate low to high relative AUCell scores, respectively.

**Extended Data Fig. 7**

**TAM gene signatures segregate into distinct similarity clusters with non-tumor disease–associated and distant metastatic macrophage programs.**

**a-c,** To assess transcriptional relationships among distinct macrophage-related gene programs, Pearson correlation coefficients were calculated between AUCell scores derived from three categories of gene sets: 19 TAM signatures, 17 non-tumor disease-associated macrophage signatures, and 20 TAM signatures from breast-cancer metastases in lung and brain. **a**, Correlation matrix between TAM and non-tumor disease-associated macrophage gene sets. **b**, Correlation matrix between TAM and metastatic breast-cancer TAM gene sets. **c**, Correlation matrix between non-tumor disease-associated macrophage gene sets and metastatic breast-cancer TAM gene sets. AUCell scores were computed from normalized single-cell RNA-seq data of macrophage subtypes. The resulting Pearson correlation matrices were visualized as heatmaps, with color intensity representing correlation coefficients (r) ranging from –1 to 1. Rows and columns were hierarchically clustered to group gene signatures with similar correlation patterns.

**Extended Data Fig. 8**

**scRNA-seq of patient-derived lung metastases identifies two macrophage subsets and G-CSF–expressing bone marrow cells colocalizing with BCCs.**

**a**, UMAP visualization of scRNA-seq data from patient-derived breast cancer lung metastasis, colored by clusters. P10_TAM1 and P10_TAM2 indicate two distinct TAM populations identified in patient 10. n = 2 mice. **b**, Dot plot showing canonical marker-gene expression for the clusters identified in (**a**). Clustering based on marker-gene profiles delineated distinct cell types. Dot color indicates scaled average expression; dot size represents the proportion of expressing cells per cluster. **c,** Representative immunofluorescence images of bone metastatic tumors stained for cancer cell (STEM121; human-specific) and G-CSF. Nuclei were counterstained with Hoechst. Scale bars, 100 μm. **d,** Pearson correlation between BCC and G-CSF fluorescence intensities (n = 183 fields). **e**, Kaplan–Meier analysis of distant metastasis-free survival in breast cancer patients (GSE7390) stratified by *G-CSFR* mRNA expression.

**Extended Data Fig. 9**

**CXCL16$^{+}$ macrophages localize exclusively within CAF-rich stromal regions.**

**a**, Representative immunofluorescence images of bone metastatic tumors from human clinical samples, stained for cancer cell (STEM121; human-specific), α-SMA, and G-CSFR. Nuclei were counterstained with Hoechst. Scale bars, 100 μm. **b**, Representative immunofluorescence images of bone metastatic tumors from human clinical samples, stained for cancer cells (STEM121), α-SMA, and macrophage markers CD68. Nuclei were counterstained with Hoechst. Scale bars, 100 μm.
